# The carnivoran adaptive landscape reveals functional trade-offs among the skull, appendicular, and axial skeleton

**DOI:** 10.1101/2024.05.06.592620

**Authors:** Chris J. Law, Leslea J. Hlusko, Z. Jack Tseng

## Abstract

Analyses of form-function relationships are widely used to understand links between morphology, ecology, and adaptation across macroevolutionary scales. However, few have investigated functional trade-offs and covariance within and between the skull, limbs, and vertebral column simultaneously. In this study, we investigated the adaptive landscape of skeletal form and function in carnivorans to test how functional trade-offs between these skeletal regions contribute to ecological adaptations and the topology of the landscape. We found that morphological proxies of function derived from carnivoran skeletal regions exhibit trade-offs and covariation across their performance surfaces, particularly in the appendicular and axial skeletons. These functional trade-offs and covariation correspond as adaptations to different adaptive landscapes when optimized by various factors including phylogeny, dietary ecology, and, in particular, locomotor mode. Lastly, we found that the topologies of the optimized adaptive landscapes and underlying performance surfaces are largely characterized as a single gradual gradient rather than as rugged, multipeak landscapes with distinct zones. Our results suggest that carnivorans may already occupy a broad adaptive zone as part of a larger mammalian adaptive landscape that masks the form and function relationships of skeletal traits.

## Introduction

The diversity of animal forms is one of the most salient patterns across the tree of life, and how morphological variation relates to the ecological diversity and survival of species across macroevolutionary time remains a core question in evolutionary biology. The varying strength of form–function relationships provide biologists with insight into the specificity of morphological structure in determining species’ abilities to carry out ecological tasks (i.e., performances), especially when behavioral observations are scarce. Thus, performance is considered the link between morphology, ecology, and fitness (Arnold 1983; Wainwright 1994; Higham et al. 2021). Functional traits are morphological, phenological, and physiological traits that affect fitness and are often used to estimate performance (Higham et al. 2021). Many researchers have examined the form-function relationship of the skull (Santana et al. 2010; Collar et al. 2014; Law et al. 2018; Tseng et al. 2023), appendicular skeleton (Sustaita et al. 2013; Dickson and Pierce 2019; Sansalone et al. 2020; Amson and Bibi 2021), and axial skeleton (Polly et al. 2016; Jones et al. 2018, 2021; Law et al. 2019; Stayton 2019), often finding trade-offs that are hypothesized to facilitate distinct ecological adaptations. For example, the gradient from short, broad jaws to long, narrow jaws is associated with a functional trade-off between generating stronger bites and quicker bites or wider gapes (Herring and Herring 1974; Dumont and Herrel 2003; Slater and Van Valkenburgh 2009; Slater et al. 2009; Forsythe and Ford 2011; Santana 2015), and similarly, the gradient from gracility to robustness in limb bones is associated with a functional trade-off between increasing cost of transport associated with cursoriality and resisting stresses associated with locomoting through resistant media (Martín-Serra et al. 2014a, 2014b; Kilbourne 2017; Hedrick et al. 2020; Muñoz 2020; Marshall et al. 2021; Rickman et al. 2023). However, most of these studies investigate trade-offs within individual bones (e.g., the mandible, humerus, or femur) and few have investigated functional trade-offs and covariation within and among the three major skeletal systems.

The rise of adaptive landscape analyses enables researchers to investigate the adaptive evolution of performance by elucidating the underlying links among morphology, ecology, and fitness benefits (i.e., adaptiveness) at the macroevolutionary level (Arnold et al. 2001). Although Ornstein–Uhlenbeck (OU) models (Hansen 1997; Butler and King 2004; Beaulieu et al. 2012; Uyeda and Harmon 2014; Bastide et al. 2018) are widely used to test for the presence of adaptive zones or peaks (e.g., Collar et al. 2014; Price and Hopkins 2015; Friedman et al. 2016; Zelditch et al. 2017; Arbour et al. 2019; Law 2022; Slater 2022), it remains difficult to characterize the full topology (i.e., peaks, valleys, and slope) of the adaptive landscape as well as assess the relative importance of multiple performance traits and their contributions to overall adaptive landscape using these models. Adaptive landscape analyses (Polly et al. 2016; Dickson and Pierce 2019; Dickson et al. 2021) can overcome these limitations by examining the distribution of species in morphospace and its relationship to the relative importance of various functional traits on the topology of the adaptive landscape. While an increasing number of studies have used functional adaptive landscapes to examine links between morphological diversity and functional performance (Polly et al. 2016; Dickson and Pierce 2019; Stayton 2019; Dickson et al. 2021; Jones et al. 2021; Tseng et al. 2023), no study has yet to investigate these relationships among the skull, limbs, and vertebral column.

Here, we examined the trade-offs and covariation among individual performance surfaces derived from functional traits of the skull, appendicular skeleton, and axial skeleton as well as assessed their relative contributions to ecological adaptations and the overall landscape. To explore these patterns, we used terrestrial carnivorans (e.g., bears, cats, dogs, weasels, and their relatives) as our model because of their high species richness and well-studied broad morphological and ecological diversity. Numerous researchers have investigated the morphological diversity of the carnivoran skull (Radinsky 1981; Van Valkenburgh 2007; Figueirido et al. 2011; Law et al. 2018, 2022; Tseng and Flynn 2018; Slater and Friscia 2019), appendicular skeleton (Van Valkenburgh 1985, 1987; Iwaniuk et al. 1999; Samuels et al. 2013; Martín-Serra et al. 2014a, 2014b), vertebral column (Randau et al. 2017; Figueirido et al. 2021; Martín-Serra et al. 2021), and overall body plan (Law 2021a, 2021b; Slater 2022). This diversity is attributed to mosaic evolution, in which different skeletal components exhibit distinct modes of evolution either from phylogenetic natural history (Uyeda et al. 2018) or from selection for ecological adaptations (Law et al. 2024). The ability of individual skeletal components to adapt to specific ecological factors independently from each other may have contributed to the clade’s hierarchical evolution. The hierarchical evolution is primarily framed by dental adaptations along an axis of dietary resource use, which are hypothesized to facilitate the early radiation of carnivorans across a rugged, multi-peak adaptive landscape (Slater and Friscia 2019). Subsequent evolution led to the continual partitioning between clades, resulting in the origination of extant carnivoran families that occupy different adaptive zones (Humphreys and Barraclough 2014) with distinct morphologies in the skull, appendicular, and axial skeletons (Law 2021a; Law et al. 2022, 2024). Skeletal variation in the mandible, hindlimb, and post-diaphragmatic region of the vertebral column then arose along shared ecological axes among taxa, theoretically leading to distinct ecological zones across the adaptive landscape (Law et al. 2022, 2024). Despite this large body of knowledge in carnivoran morphology, the functional implications of these skeletal traits remain to be tested across the adaptive landscape; that is, how do morphological traits in the skull, appendicular skeleton, and vertebral column dictate the ecological performance of carnivoran species?

Our goals of this study were three-fold. First, we described functional trade-offs and covariation among individual performance surfaces derived from functional traits from the skull, appendicular skeleton, and axial skeleton. Second, because morphological traits are often associated with locomotor and dietary adaptive peaks (Slater and Friscia 2019; Law et al. 2022, 2024; Slater 2022) and their functional trade-offs are often hypothesized to facilitate distinct ecological adaptations (Slater and Van Valkenburgh 2009; Slater et al. 2009; Martín-Serra et al. 2014b, 2014a), we tested how these performance surfaces contribute to unique adaptive landscapes and the formation of adaptive zones along locomotor, dietary, and phylogenetic axes. Third, we explored the topology of the adaptive landscape of carnivorans. Previous work using OU modeling provided evidence that morphological proxies for appendicular function exhibit relatively low ruggedness across the adaptive landscape despite also exhibiting distinct adaptive zones (Slater 2022). Adaptive landscape analyses will further clarify whether if these adaptive zones are steep peaks or broad plateaus as well as how functional traits from the skull and axial skeleton contributes to the adaptive landscape. Overall, this work provides a baseline understanding of the relative contributions of the skull, appendicular skeleton, and axial skeleton to the adaptive landscape, setting a foundation for future hypothesis testing on the processes that influence the evolution of animal form and function.

## Methods

### Morphospace and functional proxies

We created a morphospace of 109 terrestrial carnivoran species based on 136 linear and angular measurements that capture morphological variation across the entire skeleton (Fig. S1). This dataset includes seven cranial traits, seven mandibular traits, 13 forelimb traits, 13 hindlimb traits, and 11 traits each in the third cervical, fifth cervical, first thoracic, middle thoracic, diaphragmatic thoracic, last thoracic, first lumbar, middle lumbar and last lumbar vertebrae. We were able to incorporate representatives from 12 of the 13 extant terrestrial carnivoran families; specimens from Prionodontidae were unavailable. We removed size effects on linear measurements by calculating the log shape ratio (i.e., ln[trait/size]) of each skeletal trait, where size is the geometric mean of all linear trait measurements (see Sensitivity Analyses 1 in the Supplementary Materials for analyses based on non-size corrected data). We used only adult male specimens because carnivorans exhibit differing degrees of sexual dimorphism (Law 2019). We then conducted a principal component analysis (PCA) using the covariance matrix on all size-corrected measurements and used the first two PC axes (46.7% of the total variance) to create the morphospace (Fig. 1; see Table S1 for PC loadings). Carnivoran species are widely distributed across the morphospace except for the bottom-left region in which no species occupy (-PC1, -PC2). We chose not to run a phylogenetic PCA because we are interested in the primary dimensions of morphological variation regardless of phylogenetic structuring. In addition, pPCA is more difficult to interpret because it is a mixture of major axes that describe non-phylogenetic variation and scores that contain phylogenetic components of variation, and pPC axes are not orthogonal to each other, meaning that the first two axes (which we use for the adaptive landscape analyses) may include less variance explained than PCA by containing correlated variance components rather than independent ones (Polly et al. 2013).

**Fig. 1.**
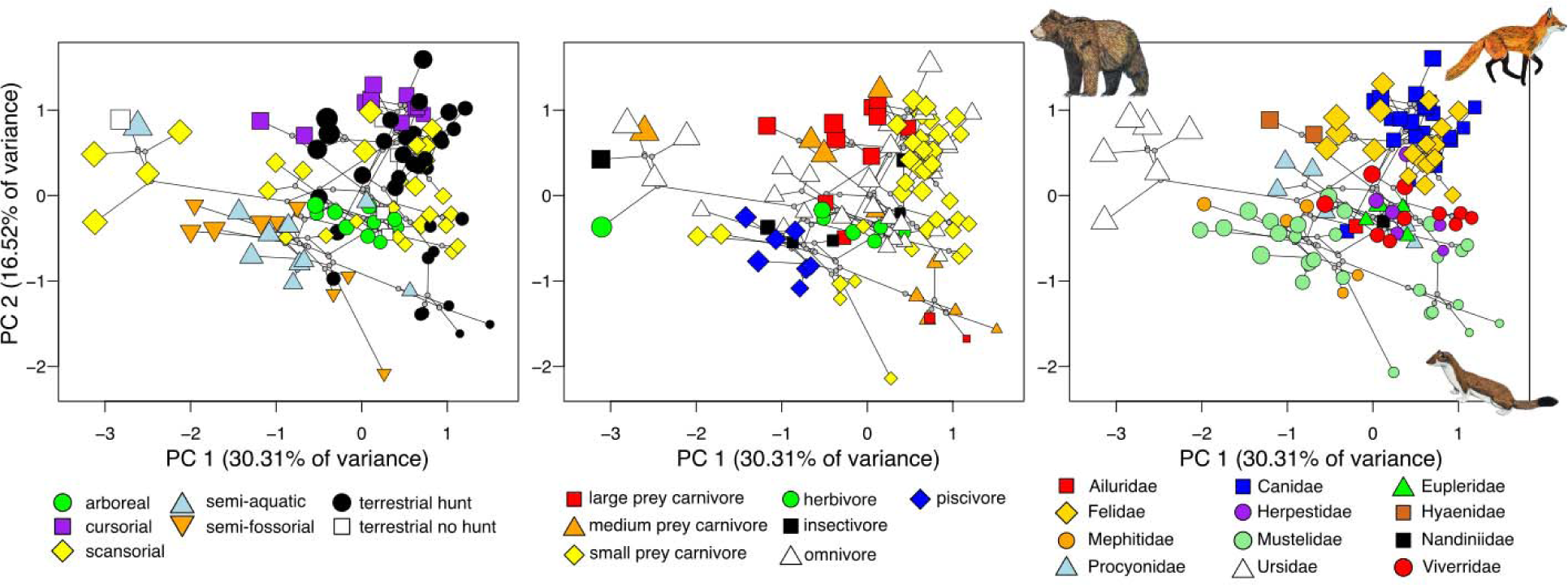
Phylomorphospace of the carnivoran skeletal system defined by principal components (PCs) 1 and 2. PCA was conducted using 136 linear and angular measurements that capture morphological variation across the skull, appendicular, and axial skeletons (Table S1 shows trait loadings). Linear measurements were size-corrected using log shape ratios. Size of points are scaled to estimated body size based on the geometric mean of all measurements.

From the 136 morphological traits, we then calculated 27 morphological proxies of function as proxies for functional traits (hereinafter called “functional proxies”; Table 1; see Supplementary Materials for full biomechanical and ecomorphological justification). These functional proxies are often used to capture the functional diversity of the skull (Greaves 2012), limbs (Davis 1964; Samuels et al. 2013), and vertebral column (Boszczyk et al. 2001; Jones et al. 2020, 2021).

**Table 1.**
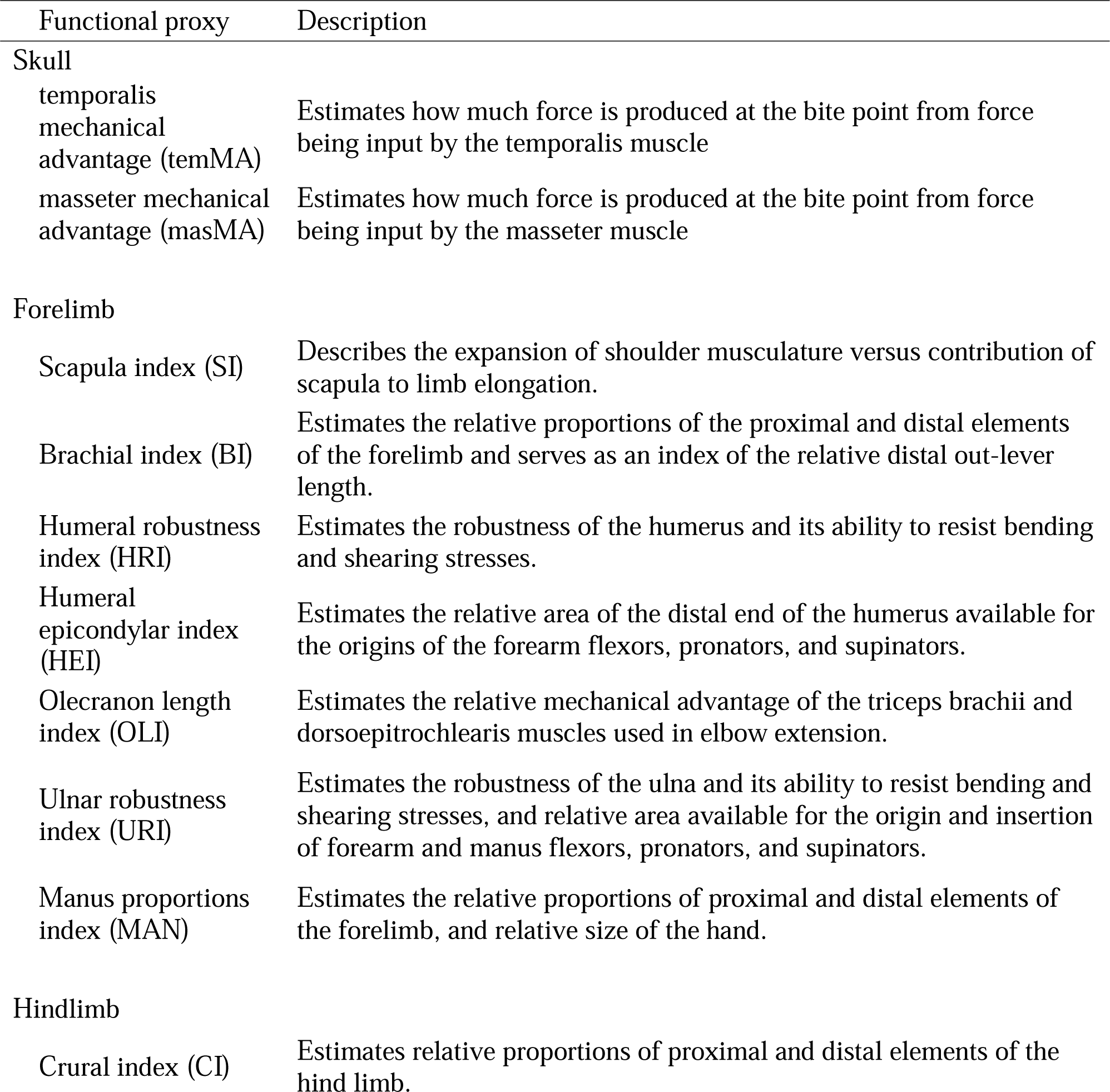
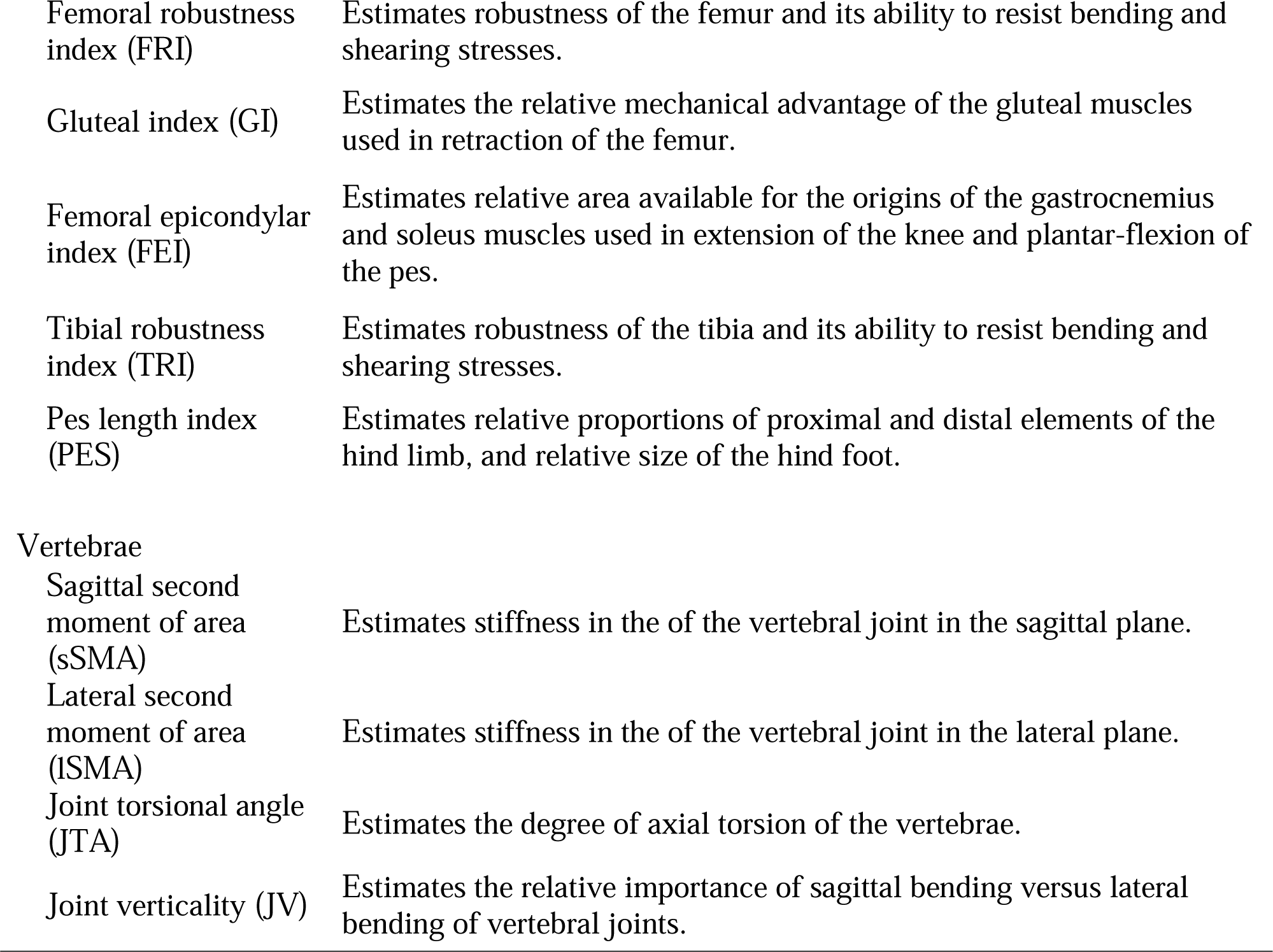
Functional proxies capturing the functional diversity of the skull (Greaves 2012), limbs (Davis 1964; Samuels et al. 2013), and vertebral column (Boszczyk et al. 2001; Jones et al. 2020, 2021). The full performance, biomechanical, and/or ecomorphological justification for each functional proxy is expanded upon in the Supplementary Materials.

| Functional proxy | Description |
| --- | --- |
| Skull |  |
| temporalis mechanical advantage (temMA) | Estimates how much force is produced at the bite point from force being input by the temporalis muscle |
| masseter mechanical advantage (masMA) | Estimates how much force is produced at the bite point from force being input by the masseter muscle |
| Forelimb |  |
| Scapula index (SI) | Describes the expansion of shoulder musculature versus contribution of scapula to limb elongation. |
| Brachial index (BI) | Estimates the relative proportions of the proximal and distal elements of the forelimb and serves as an index of the relative distal out-lever length. |
| Humeral robustness index (HRI) | Estimates the robustness of the humerus and its ability to resist bending and shearing stresses. |
| Humeral epicondylar index (HEI) | Estimates the relative area of the distal end of the humerus available for the origins of the forearm flexors, pronators, and supinators. |
| Olecranon length index (OLI) | Estimates the relative mechanical advantage of the triceps brachii and dorsoepitrochlearis muscles used in elbow extension. |
| Ulnar robustness index (URI) | Estimates the robustness of the ulna and its ability to resist bending and shearing stresses, and relative area available for the origin and insertion of forearm and manus flexors, pronators, and supinators. |
| Manus proportions index (MAN) | Estimates the relative proportions of proximal and distal elements of the forelimb, and relative size of the hand. |
| Hindlimb |  |
| Crural index (CI) | Estimates relative proportions of proximal and distal elements of the hind limb. |
| Femoral robustness index (FRI) | Estimates robustness of the femur and its ability to resist bending and shearing stresses. |
| Gluteal index (GI) | Estimates the relative mechanical advantage of the gluteal muscles used in retraction of the femur. |
| Femoral epicondylar index (FEI) | Estimates relative area available for the origins of the gastrocnemius and soleus muscles used in extension of the knee and plantar-flexion of the pes. |
| Tibial robustness index (TRI) | Estimates robustness of the tibia and its ability to resist bending and shearing stresses. |
| Pes length index (PES) | Estimates relative proportions of proximal and distal elements of the hind limb, and relative size of the hind foot. |
| Vertebrae |  |
| Sagittal second moment of area (sSMA) | Estimates stiffness in the of the vertebral joint in the sagittal plane. |
| Lateral second moment of area (lSMA) | Estimates stiffness in the of the vertebral joint in the lateral plane. |
| Joint torsional angle (JTA) | Estimates the degree of axial torsion of the vertebrae. |
| Joint verticality (JV) | Estimates the relative importance of sagittal bending versus lateral bending of vertebral joints. |

### Ecological traits

We classified the 109 carnivoran species into one of seven locomotor regimes: arboreal (species that primarily live and forage in trees and rarely come down to the ground), cursorial (species that displays rapid bounding locomotion, particularly during hunting), scansorial (species that spend equal time in trees and on the ground), semi-aquatic (species that regularly swim for dispersal and/or foraging), semi-fossorial (species that regularly dig for shelter and/or foraging), and terrestrial (species that primarily live on the ground and rarely run, climb, dig, or swim during foraging). Terrestrial species were further categorized as terrestrial hunters (species that exhibit ambush and/or pouncing behaviors to kill prey) and terrestrial non-hunters (species that rarely hunt for prey). We also classified each species into one of seven dietary regimes: large prey hypercarnivory (consist of >70% terrestrial vertebrate prey that exceeds the predator’s own body mass), medium prey hypercarnivory (consist of >70% terrestrial vertebrate prey that are up to the predator’s own body mass), small prey hypercarnivory (consist of >70% terrestrial vertebrate prey that are up to 20% of the predator’s own body mass), omnivory (consist of >50% terrestrial vertebrates), insectivory (consisting of >70% invertebrates), aquatic carnivory (consist of >90% aquatic prey), and herbivory (consist of >90% plant material). These locomotor and dietary regimes are widely used to describe carnivoran ecology and have demonstrated significant associations with various traits of the cranial, appendicular, and axial skeletons of carnivorans (Van Valkenburgh 1987, 2007; Friscia et al. 2007; Samuels et al. 2013). Categorization for locomotor and dietary regimes were obtained from previous work (Van Valkenburgh 1985; Samuels et al. 2013; Law 2021a) with minor edits based on literature review.

To check the relationship between the morphospace (i.e., PCs 1 and 2) with ecological regimes, we conducted two multivariate ANOVA models (i.e., morphospace ∼ locomotor mode and morphospace ∼ dietary ecology) in the R package RRPP v1.4.0 (Adams and Collyer 2018). We found that the morphospace exhibited a significant relationship with locomotor mode (R^2^ = 0.29, F = 6.78, P = 0.001) and with dietary ecology (R^2^ = 0.16, F = 3.12, P = 0.001). Post-hoc pairwise comparison tests also indicated significance differences among most ecological regimes (Table S2). Testing using multivariate PGLS models (Clavel et al. 2019; Clavel and Morlon 2020) also indicated that the exhibited a significant relationship with locomotor mode (Pagel’s λ = 0.97, Pillai’s trace = 0.202, P = 0.041) and with dietary ecology (Pagel’s λ = 0.97, Pillai’s trace = 0.213, P = 0.017).

### Performance surfaces and adaptive landscapes

We investigated the functional optimality of the skeleton using adaptive landscape analyses (Polly et al. 2016; Dickson and Pierce 2019; Dickson et al. 2021; Jones et al. 2021) in the R package Morphoscape (Dickson et al. 2021). We first created 27 performance surfaces by interpolating each of the 27 functional proxies across the morphospace surface using ordinary kriging. Initial inspection of these surfaces revealed that the performance surfaces of four functional proxies (masseter mechanical advantage [masMA], scapula index [SI], gluteal index [GI], and tibial robustness [TRI]) exhibited topological peaks and valleys that outlined clusters of species and even single species (Fig. 2; see section below about the downsides to using empirical data instead of theoretical data). Therefore, we removed these four functional proxies from subsequent analyses.

**Fig. 2.**
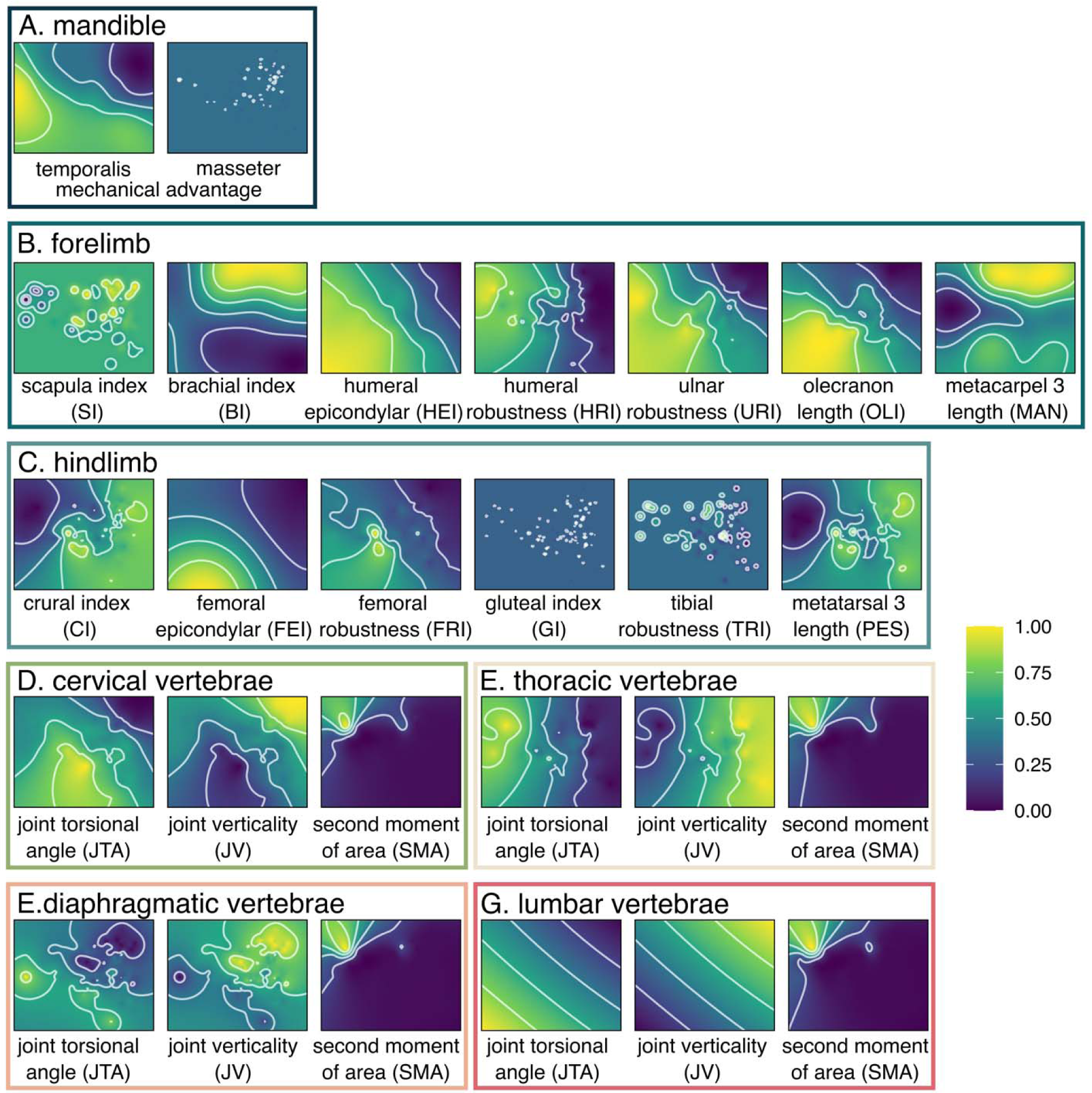
Performance surfaces for each functional proxy. Color represents height on the performance surface. See Table 1 for definitions of functional proxies. We removed masMA, SI, GI, and TRI in subsequent adaptive landscape analyses.

We computed a combined adaptive landscape (W) as the summation of all 23 performance surfaces (F_n_), each weighted by their relative importance or contribution to overall fitness (w_n_) (Polly et al. 2016):

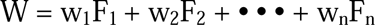

where W is optimized as the likelihood of combinations of performance surface and relative fitness, under the definition that the total fitness sums to 1 and the variance of all surfaces is equal (Polly et al. 2016). We tested all possible combinations of weights, ranging from 0 to 1 in increments of 0.25, across a total of 27,405 possible adaptive landscapes (see Sensitivity Analyses 2 in the Supplementary Materials for further investigation of the concern with number of partition weight increments).

We identified the optimally weighted landscape that maximizes the fitness of each locomotor regime using the function calcWprimeBy and tested whether these optimal landscapes are significantly different among locomotor ecological groups using the function multi.lands.grp.test. Significance testing for differences among landscapes was performed by comparing the number of landscapes shared by the top 5% of each group with the total number of landscapes in the top 5% of models (Jones et al. 2021). The top percentile of each group was determined using a chi-squared test. We also investigated differences in adaptive landscapes among dietary groups and carnivoran families that contained more than one species.

### Creating adaptive landscapes using theoretical traits

A great concern in using empirical data in creating performance surfaces and adaptive landscapes is that empirical-specific values are always less evenly distributed in the morphospace. The unevenness contributes to heterogeneous resolution of the interpolation applied to the space by the ordinary kriging method, and thus unevenness in the landscape itself. Denser sampled regions will be more likely to have topological relief (i.e., peaks and valleys) than sparsely sampled regions when using actual specimen values, not necessarily because of a real underlying peak there. To mitigate these issues, many researchers have used theoretical data across an evenly spaced grid to create performance surfaces and adaptive landscapes (Polly et al. 2016; Dickson and Pierce 2019; Smith et al. 2021; Tseng et al. 2023; Sansalone et al. 2024).

Therefore, we investigated whether there is consistency in our adaptive landscape analyses when using functional proxies derived from actual species versus using functional proxies derived from theoretical species. To fully sample skeletal variation throughout the morphospace, we generated 63 theoretical species evenly across the morphospace in a 9 x 7 grid along the first two PCs. We then generated theoretical morphological traits from each of the 63 theoretical species and calculated the 27 functional proxies (Table 1). We then performed the same procedures as described above to generate the 27 performance landscapes and adaptive landscapes optimized for locomotor ecology, diet, and family. We found that the patterns found in adaptive landscapes are similar with only slight differences. To avoid confusion between the two approaches, we report the results using the theoretical morphologies in the Supplementary Results of the Supplementary Materials.

### Sensitivity Analyses

Lastly, we conducted sensitivity analyses to investigate whether (1) removing the effects of size on the morphological traits in creating the morphospace and (2) the coarse increments of partition weights may potentially influence our interpretation of the results. We found that these factors do not affect our findings (see Sensitivity Analyses section in the Supplementary Materials).

## Results

### Performance surfaces reveal trade-offs and covariation within skeletal systems

Each functional proxy mapped onto the morphospace revealed both unique and similar performance surfaces that characterize trait groups, suggesting that functional trade-offs and covariation are present within the skull, appendicular skeleton, and axial skeleton. In the skull, mechanical advantage of the temporalis (temMA) highest towards the left and bottom-left (-PC1, -PC2) of morphospace and declining towards the top-right (+PC1, +PC2). In contrast, there is no distinct pattern in masseter mechanical advantage (masMA) (Fig. 2).

In the forelimb, there is a trade-off between limb elongation and elbow robustness: functional proxies of radius (BI) and metacarpal (MAN) elongation are highest in the top-right (+PC1, +PC2) of morphospace. BI transitions towards increased robustness to the bottom-right (+PC1, -PC2), whereas MAN transitions towards increased robustness to the left (-PC1). Proxies associated with increased mechanical advantage of elbow extension (OLI) and attachment sites for forearm flexor, pronator, and supinator muscles on the humeral epicondyles (HEI) and ulna (URI) are highest in the bottom-left and lowest in the top-right. Overall robustness of humerus (HRI) is highest on the left side of the morphospace (-PC1) and transitions towards increased elongation to the right (+PC1), following a similar distribution as the latter indices. There is no distinct pattern in scapula index (SI). The hindlimb also exhibits a trade-off between elongation and robustness: indices of tibial (CI) and metatarsal (PES) elongation tend to be highest in the right side (+PC1, with CI also trending towards -PC2) and transitions to increased robustness towards to the left side (-PC2), whereas indices for femoral (FEI, FRI) robustness tend to be highest on the bottom (-PC2) but transitions towards less robustness in the top-right (+PC1, +PC2). There are no distinct patterns in indices for gluteal muscles (GI) and tibial robustness (TRI).

In the vertebral joints, the performance surfaces show a trade-off between joint torsional angle (JTA) as a proxy for range of rotational motion and joint verticality (JV) as a proxy for sagittal mobility. For the cervical, diaphragmatic, and lumbar joints, JTA is highest in the bottom-left of the morphospace (-PC1, -PC2) and declines diagonally to the top-right (+PC1, +PC2), whereas JV exhibits the opposite pattern (i.e., highest in the top-right and lowest in the bottom-left). The thoracic vertebra exhibits similar JTA and JV distributions but in the horizontal plane (i.e., highest JTA in the left side of morphospace and highest JV in the right). In all vertebral joints, second moment of area (SMA) as a proxy for stiffness tends to be greatest towards the top-left of morphospace (-PC1, +PC2) and declines towards the right side of morphospace (+PC1).

### Optimized adaptive landscapes reveal trade-offs and covariation among skeletal systems

After summation of all performance surfaces based on optimized weights, we found that the combined adaptive landscape is heavily weighted for sagittal mobility of the pre-diaphragmatic thoracic (w_JV_ = 0.41) and lumbar (w_JV_ = 0.38) regions (Fig. 3A; Table S3). When adaptive landscapes are optimized by locomotor ecologies, we found different degrees to which the 23 functional proxies are incorporated among the different adaptive landscapes. The cursorial landscape is characterized by lengthening of the forelimb, particularly in the metacarpal (w_MAN_ = 0.56) and, to a lesser extent, the radius (w_BI_ = 0.04) (Fig. 3C; Table S3). The cursorial landscape is also strongly weighted with functional proxy associated with increased sagittal mobility of the pre-diaphragmatic thoracic joints (w_JV_ = 0.38). The semi-aquatic and semi-fossorial landscapes do not significantly differ from one another (P = 0.184; Table 2), and both are similarly weighted for larger humeral epicondyles (semi-aquatic w_HEI_ = 0.28; semi-fossorial w_HEI_ = 0.40) and increased joint torsion in the cervical joints (semi-aquatic w_JTA_ = 0.39; semi-fossorial w_JTA_ = 0.11) (Fig. 3E, F; Table S3). The semi-aquatic landscape is further strongly weighted for more robust ulna (w_URI_ = 0.16), whereas the semi-fossorial landscape is further strongly weighted for increased joint torsion in the diaphragmatic joint (w_JTA_ = 0.37) (Fig. 3E, F; Table S3). The terrestrial non-hunter landscape is not significantly different from the cursorial and semi-fossorial landscapes (Table 2). This landscape is strongly weighted for increased joint torsion in the diaphragmatic joint (w_JTA_ = 0.60) and lengthening of the metacarpal (w_MAN_ = 0.21) (Fig. 3H; Table S3). The remaining locomotor landscapes (i.e., arboreal, scansorial, and terrestrial hunter) do not significantly differ from one another (Table 2) and are heavily weighted for sagittal mobility of the pre-diaphragmatic thoracic and lumbar joints (Fig. 3B, D, G; Table S3).

**Fig. 3.**
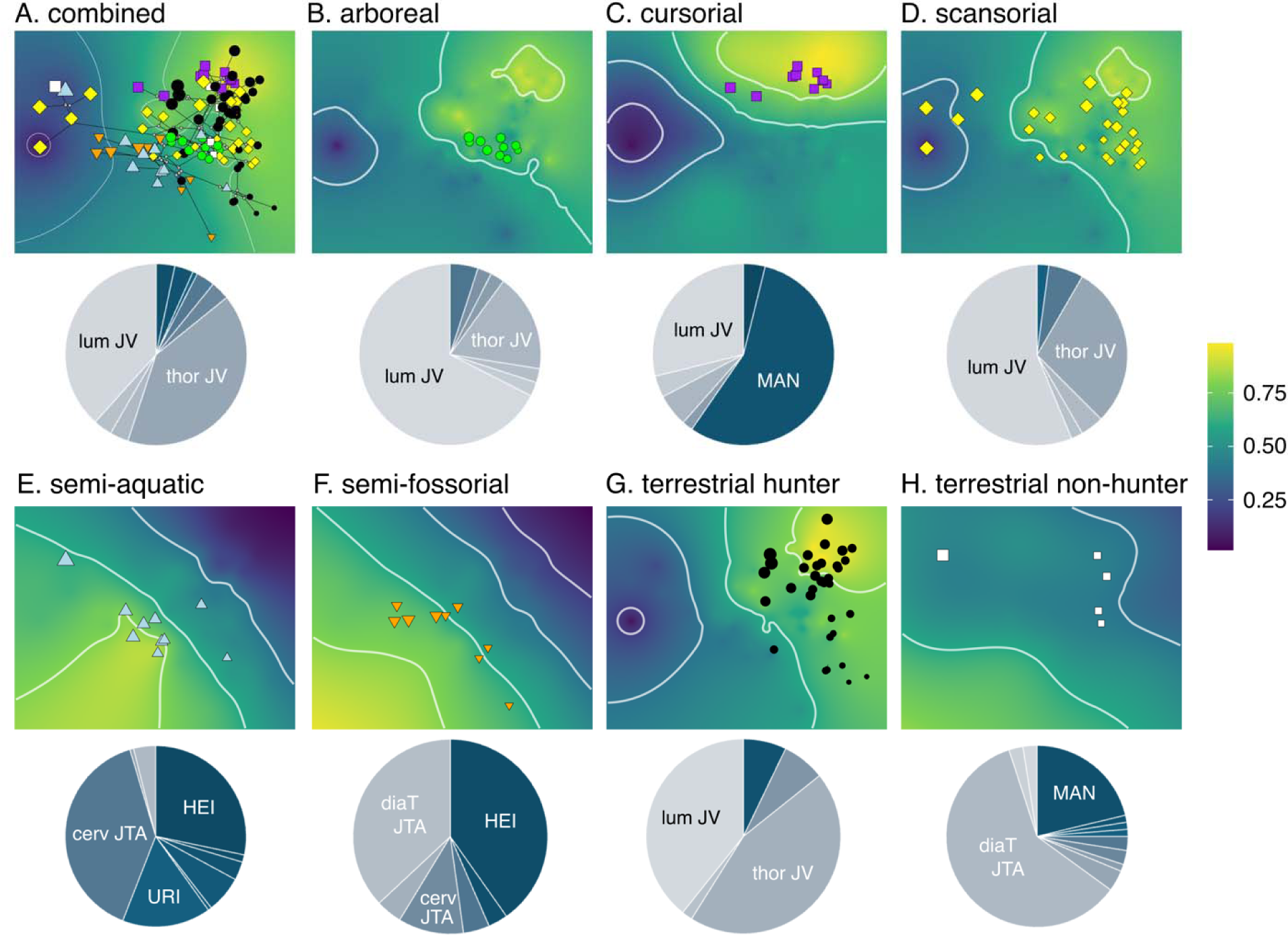
Adaptive landscapes optimized for all carnivorans and each locomotor group. Landscapes were produced by combining the performance surfaces and optimizing their weightings to maximize the height of the landscape at the group mean. Pie charts show the relative weights of each performance surface on each landscape (Table S3 shows breakdown of weights). Functional proxies with weights > 0.07 were labeled. Table 2 shows statistical tests comparing adaptive landscapes among locomotor groups. Size of points are scaled to estimated body size based on the geometric mean of all measurements.

**Table 2.** Pairwise significance tests among locomotor adaptive landscapes. Top triangle: number of landscape models shared in the top 5% between the paired groups. Bottom triangle: P-values for difference between groups. Bolded P-values indicate significance.

|  | arboreal | cursorial | scansorial | semi-aquatic | semi-fossorial | terrestrial hunter | terrestrial non-hunter |
| --- | --- | --- | --- | --- | --- | --- | --- |
| arboreal | - | 4 | 7 | 0 | 0 | 6 | 6 |
| cursorial | 0.100 | - | 10 | 0 | 0 | 7 | 34 |
| scansorial | 0.800 | 0.077 | - | 0 | 0 | 14 | 15 |
| semi-aquatic | **0.001** | **0.001** | **0.001** | - | 88 | 0 | 0 |
| semi-fossorial | **0.001** | **0.001** | **0.001** | 0.184 | - | 0 | 0 |
| terrestrial hunter | 0.500 | 0.231 | 0.417 | **0.001** | **0.001** | - | 12 |
| terrestrial nonhunter | **0.001** | 0.077 | **0.001** | **0.001** | 0.174 | **0.001** | - |

When adaptive landscapes are optimized by dietary ecologies, we found that significant differences in landscapes appear associated with piscivory (Fig. S2; Table S4). The piscivorous adaptive landscape is similarly weighted for larger humeral epicondyles (w_HEI_ = 0.32), more robust ulna (w_URI_ = 0.22), and increased joint torsion in the cervical joints (w_JTA_ = 0.22) (Fig. S2; Table S5). In contrast, adaptive landscapes based on other diets are not significantly different from each other and are largely characterized by increased sagittal mobility of the pre-diaphragmatic thoracic and/or lumbar joints (Fig. S2; Table S4; Table S5).

Lastly, we found different adaptive landscapes when they are optimized by family. Adaptive landscapes for felids, viverrids, euplerids, herpestids, canids, and procyonids are not significantly different from each other and all remain heavily weighted for sagittal mobility of the pre-diaphragmatic thoracic and/or lumbar joints (Fig. 4; Table S6; Table S7). The canid landscape is also weighted for lengthening of the metacarpal (Fig. 4G; Table S6). The mephitid and mustelid landscapes resemble semi-fossorial and semi-aquatic landscapes, respectively. The mephitid landscape is equally weighted by larger humeral epicondyles (w_HEI_ = 0.38) and increased joint torsion in the diaphragmatic joint (w_JTA_ = 0.38) (Fig. 4I; Table S6), whereas the mustelid landscape is weighted by increased sagittal mobility of the pre-diaphragmatic thoracic joints (w_JV_ = 0.22), more robust ulna (w_URI_ = 0.17), larger humeral epicondyles (w_HEI_ = 0.13), and increased joint torsion in the cervical (w_JTA_ = 0.18) and diaphragmatic (w_JTA_ = 0.13) joints (Fig. 4K; Table S6). Only the mephitid landscape significantly differs from all other families (Table S7). The hyaenid landscape weighted heavily for elongation of the radius (w_BI_ = 0.50) and metacarpal (w_MAN_ = 0.50) (Fig. 4D). Lastly, the ursid landscape weighted heavily for larger humeral epicondyles (w_HEI_ = 0.38), increased joint torsion in the thoracic joints (∑w_JTA_ = 0.41), and increased robustness of the ulna (w_URI_ = 0.14) and humerus (w_HRI_ = 0.08) (Fig. 4H). Both hyaenid and ursid landscapes significantly differ with most other family-specific landscapes (Table S7).

**Fig. 4.**
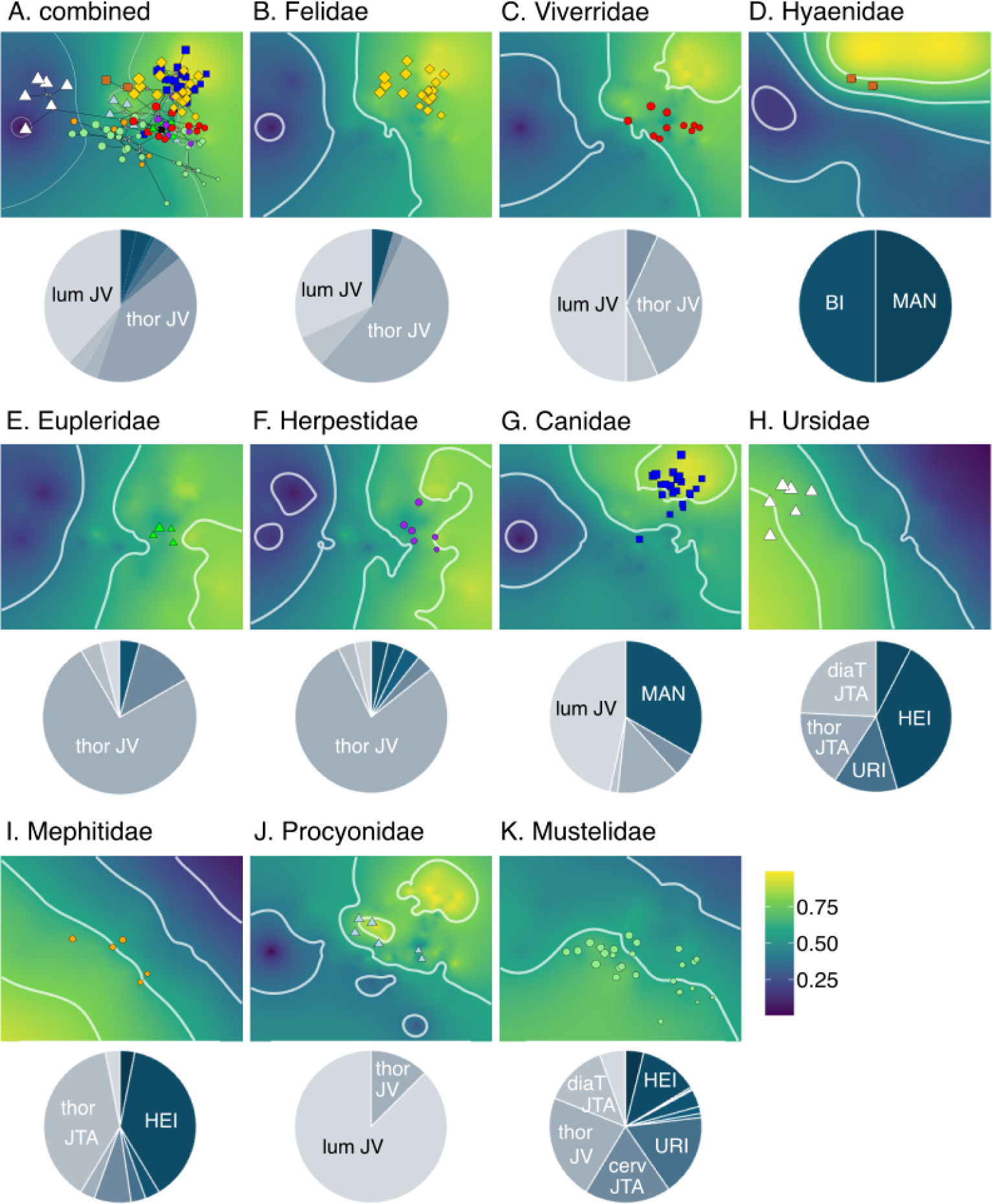
Adaptive landscapes optimized for all carnivorans and each family. Landscapes were produced by combining the performance surfaces and optimizing their weightings to maximize the height of the landscape at the group mean. Pie charts show the relative weights of each performance surface on each adaptive landscape (Table S6 shows breakdown of weights). Functional proxies with weights > 0.09 were labeled. Table S7 shows statistical tests comparing adaptive landscapes among families. Size of points are scaled to estimated body size based on the geometric mean of all measurements.

## Discussion

The diversity found in the carnivoran skeletal system is attributed to mosaic evolution, in which the mandible, hindlimb, and post-diaphragmatic vertebrae showed evidence of adaptation towards ecological regimes whereas the cranium, forelimb, and pre-diaphragmatic vertebrae reflect clade-specific evolutionary shifts (Law et al. 2024). Using adaptive landscape analyses, we further found that functional proxies derived from this morphological diversity exhibit trade-offs and covariation, particularly within and between the appendicular and axial skeletons. These functional trade-offs and covariation corresponded as adaptations to different adaptive landscapes when optimized by various factors including phylogeny, dietary ecology, and, in particular, locomotor mode. Lastly, these adaptive landscapes and underlying performance surfaces were characterized by rather broad slopes, hinting that carnivorans occupy a broad adaptive zone as part of a larger mammalian adaptive landscape that masks the form and function relationships of skeletal traits.

### Performance surfaces reveal trade-offs and covariation among skeletal systems

In the appendicular skeleton, we found support that the gradient from gracility to robustness often found as the primary source of variation across limb bone morphospace (Martín-Serra et al. 2014a, 2014b; Kilbourne 2017; Hedrick et al. 2020; Muñoz 2020; Marshall et al. 2021; Rickman et al. 2023) signifies a functional trade-off between increasing cost of transport associated with cursoriality and resisting stresses associated with locomoting through resistant media (Fig. 2B). Specifically, long, gracile limb bones particularly on the distal ends of the limbs facilitate increased stride length and decreased moment of inertia of limbs, which in turn decreases the energetic cost of transport and increases running speeds (Kram and Taylor 1990; Strang and Steudel 1990; Garland and Janis 1993; Polly 2007; Pontzer 2007a, 2007b; Kilbourne and Hoffman 2013). In contrast, short, robust limb bones facilitate resistance to bending and shearing stresses and increased mechanical advantage for forceful movements by reducing the out-lever of the limb and increasing the in-lever of muscle forces (Hildebrand 1985a; Nakai and Fujiwara 2023). Robustness also permits increased surface area of the bone for more muscles to attach. Enlargement of the humeral and femoral epicondyles increases the attachment sites of several muscles (i.e., flexors, pronators, and supinators in the forelimb and gastrocnemius and soleus muscles in hindlimb) responsible for generating power, force, and stability (Davis 1964; Hildebrand 1985a; Lessa and Stein 1992; Lagaria and Youlatos 2006). Additionally, an enlarged olecranon process facilitates stronger extension and flexion of the elbow and wrist by increasing mechanical advantage of the triceps brachii and dorsoepitrochlearis muscles and providing greater attachment sites for the ulnar head of the flexor carpi ulnaris (Davis 1964; Hildebrand 1985a; Lessa and Stein 1992; Lagaria and Youlatos 2006). These adaptations facilitate the ability to generate large forces during certain locomotor behaviors such as digging (Hildebrand 1985a; Lessa and Stein 1992; Lagaria and Youlatos 2006; Samuels et al. 2013; Rose et al. 2014; Rickman et al. 2023) or swimming (Fish 2000; Samuels and Van Valkenburgh 2008; Samuels et al. 2013; Kilbourne 2017).

In the axial skeleton, our investigation using functional proxies suggested that carnivorans exhibit trade-offs between joint mobility and range of axial rotation. That is, high sagittal mobility covaries with low range of axial rotation whereas low sagittal mobility covaries with high range of axial rotation. This pattern was surprising because, compared to other tetrapods, mammals exhibit intervertebral joints that are characterized by high sagittal mobility and high axial rotation (Jones et al. 2021). A possible explanation was that the covariation between high sagittal mobility and low axial rotation may serve as a further adaptation to increasing forward locomotion by prioritizing flexibility in the sagittal plane through the reduction of torsional twisting. High mobility of the backbone in the sagittal plane has long been recognized as a key adaptation facilitating the diversity of different locomotor habits in mammals, particularly asymmetrical gaits (e.g., gallop, half-bound, bound) by enabling extensive dorsoventral flexion of the body axis (Hildebrand 1959, 1985b; Gambaryan 1974; Schilling and Hackert 2006). The reduction of torsional twisting in carnivorans may, therefore, prioritize force generation in the sagittal plane (rather than parasagittal or transverse plane) needed for these locomotor behaviors. Evidence for this hypothesis was found in comparisons with ungulates, where carnivorans exhibit up to ∼38° more sagittal mobility in the lumbar region and up to 200% less axial rotational mobility in the thoracic region compared to ungulates (Belyaev et al. 2021, 2022, 2023). Increased rotational mobility of the backbone in ungulates is hypothesized to enhance agile maneuvering such as sharp cornering and quick directional changes when escaping from predators (Belyaev et al. 2023).

### Functional covariation between appendicular and axial skeletons optimizes the adaptive landscapes of some locomotor ecologies

The optimized adaptive landscape is heavily weighted for sagittal mobility of the pre-diaphragmatic thoracic and lumbar joints (Fig. 3A), indicating that flexibility in the sagittal plane serves an important functional role for all carnivorans. When the adaptive landscapes are optimized based on locomotor mode, diet, or family, we found that locomotor behavior could provide an explanation for most landscape patterns. In our analyses of locomotor landscapes, we found that semi-aquatic and semi-fossorial landscapes were not significantly different to each other but are distinct to other locomotor landscapes (Fig. 3; Table 2). The peaks of both semi-aquatic and semi-fossorial landscapes occur in the bottom-left regions of morphospace, and species with either locomotor modes occur in overlapping regions of the landscapes. In contrast, the peaks of most of the remaining locomotor landscapes, particularly the cursorial and terrestrial hunter landscapes, occur near the top-right region of morphospace. The opposing locations of these landscape peaks corresponds to the functional trade-offs identified by the performance surfaces, suggesting that covariation of the appendicular and axial skeletons facilitate adaptations to each of these locomotor behaviors at these extreme ends. These functional trade-offs are largely independent of size effects because body size variation scales from the top-left to bottom-right of morphospace whereas the trade-offs scale from the top-right to the bottom-left (Fig. 3; Fig. 4; Fig. S2). Sensitivity analyses examining the performance surfaces and adaptive landscapes using non-size corrected morphological traits confirm this pattern (see Supplementary Materials).

Covariation of the appendicular and axial skeletons and its role in facilitating adaptations to locomotor behaviors is apparent in semi-aquatic and semi-fossorial species. Both behaviors require large force generation for swimming and digging, and the appendicular and axial skeletons of semi-aquatic and semi-fossorial carnivorans are functionally adapted for increased elbow extension through enlarged elements of the limbs and increased axial rotation in the vertebral column (Fig. 3E, F; Table S3). It is well documented that adaptations in the elbows and knees facilitate the ability to generate large power strokes for turning and stabilizing the body while swimming (Fish 2000; Samuels and Van Valkenburgh 2008; Samuels et al. 2013; Kilbourne 2017). These similar adaptations also enable semi-fossorial species to generate large forces to dig (Hildebrand 1985a; Lessa and Stein 1992; Lagaria and Youlatos 2006; Samuels et al. 2013; Rose et al. 2014; Rickman et al. 2023) and improve stability and load transfer during clearing (Hildebrand 1985a; Casinos et al. 1993; Samuels and Van Valkenburgh 2008; Rickman et al. 2023). What remains largely undetermined is the importance of increased axial joint rotation during swimming or digging. Presumably, for swimmers, torsional rotation of the intervertebral joints increases the maneuverability and ability to perform rapid turns in water (Fish 1994; Fish et al. 2003). Fully aquatic seals exhibit more flexible and compliant intervertebral joints compared to terrestrial mammals (Gál 1993); whether their joints are also more capable for torsional rotation remains to be studied. For semi-fossorial carnivorans, increased axial rotation of the vertebral column may provide additional leverage when digging through sediment. Evidence for increased axial rotation has been observed in the semi-fossorial nine-banded armadillo; experimentation on intervertebral joint flexion in this species revealed rotational motion in the joints despite not being explicitly tested (Oliver et al. 2016). Nonetheless, the benefits of increased joint rotation for digging remains puzzling. Interestingly, no carnivorans, including semi-aquatic and semi-fossorial species, occupied the highest regions (bottom-left) of semi-aquatic or semi-fossorial landscapes. A likely explanation was that further axial rotation is biologically impossible for these carnivorans given their vertebral morphology that may be under evolutionary constraints having originated from terrestrial carnivorans. Thus, semi-aquatic and semi-fossorial carnivorans may already be at the highest region of the adaptive landscape that is biologically feasible.

Cursorial species tend to be most concentrated in the top-right region of morphospace, and thus appears to serve as the opposing extreme to the semi-aquatic and semi-fossorial landscapes. The majority of cursorial carnivorans occupy regions of morphospace that corresponded to the highest regions of the cursorial landscape. This landscape indicates that the appendicular and axial skeletons of cursorial carnivorans are functionally adapted for increased stride length through elongation of forelimb and increased sagittal flexibility of the full vertebral column (Fig. 3C; Table S3). As described above, these adaptations increase running speeds and reduce the energetic cost of transport by prioritizing dorsoventral flexion and extension in the sagittal plane (Hildebrand 1959, 1985b; Gambaryan 1974; Kram and Taylor 1990; Strang and Steudel 1990; Garland and Janis 1993; Schilling and Hackert 2006; Pontzer 2007a, 2007b; Kilbourne and Hoffman 2013; Belyaev et al. 2023).

The remaining locomotor landscapes were heavily weighted for sagittal mobility of the pre-diaphragmatic thoracic and/or lumbar joints (Fig. 3; Table S3). That the arboreal landscape was not heavily weighted by additional functional proxies is surprising considering that arboreality is often described as a specialized form of locomotion (Young 2023). A possible explanation was that carnivorans do not display the full diversity of arboreal behaviors (e.g., brachiation, leaping, suspensory climbing) performed by other mammals. Another possibility was that we did not include all possible functional proxies in our analyses. For example, the ratio between proximal manual phalanx length and metacarpal length has been shown to accurately predict climbing frequency in rodents (Nations et al. 2019), and thus may have altered the optimized weights of the arboreal adaptive landscape if this proxy or others were included in this current study. These unaccounted sources may also explain why arboreal species lie away from the highest regions of the adaptive landscape (Fig. 3B).

Adaptive landscapes optimized based on family also demonstrated similar patterns as locomotor-specific landscapes (Fig. 4; Table S6). Most family-specific landscapes were heavily weighted for sagittal mobility of the pre-diaphragmatic thoracic and/or lumbar joints. Canids, which primarily exhibit cursorial or terrestrial hunting behaviors, exhibited similar patterns with the cursorial landscape with increased sagittal mobility in other regions of the vertebral column and elongation of the metacarpal (Fig. 4G; Table S6). Likewise, mephitids and mustelids comprise of many semi-aquatic and semi-fossorial species and thus exhibited similar patterns with the semi-aquatic and semi-fossorial landscapes of increased intervertebral joint rotation and enlarged limb joints (Fig. 4I, K). Adaptive landscapes for hyaenids and ursids were both unique compared to other family-specific landscapes (Table S6). The hyaenid landscape was heavily weighted for relative elongation of the forelimb (Fig. 4D); however, a caveat was that our low sample size of just two species reflected a biased representation of taxa with elongate forelimb and sloped back found in their extant diversity. Lastly, the ursid landscape was heavily weighted for increased robustness of the stylopodia of the limbs (Fig. 4H). These results were unsurprising as ursids are the biggest terrestrial carnivorans and these traits support their heavy bodies against the effects of gravity (Polly 2007; Jones et al. 2021).

Lastly, we found that only the piscivorous landscape was significantly different from all other landscapes optimized based on dietary ecology (Table S4). Unsurprisingly, the piscivorous landscape resembles the semi-aquatic landscape and was heavily weighted for larger humeral femoral epicondyles and more robust ulna as well as increased joint torsion in the cervical joints (Fig. S2; Table S5). Similarly, the insectivorous landscape resembles the semi-fossorial landscape with heavy weights towards increased joint torsion and more robust forelimbs. These results demonstrate the adaptations facilitate not only locomotor behaviors such as swimming and digging but also dietary ecologies in concert; that is, they need to swim or dig for their prey. Nevertheless, our functional proxies for feeding consisted of just the mechanical advantage of jaw closure and thus may not capture the full functional diversity found in carnivorans. The mammalian skull contains many functional trade-offs such as among bite strength, bite velocity, and gape size (Herring and Herring 1974; Dumont and Herrel 2003; Slater and Van Valkenburgh 2009; Slater et al. 2009; Forsythe and Ford 2011; Santana 2015). Thus, inclusion of additional functional proxies from the cranium, mandible, and dentition may uncover further important contributions of the skull in the evolution of carnivorans (Tseng and Flynn 2018; Slater and Friscia 2019; Law et al. 2022; Sansalone et al. 2024).

Although biomechanical and ecomorphological studies have linked many of our selected functional proxies with performance traits (Davis 1964; Boszczyk et al. 2001; Greaves 2012; Samuels et al. 2013; Jones et al. 2020, 2021), we acknowledge that our analyses were based on morphological proxies of function rather than empirical performance traits. These may affect the findings presented in this study. Future work incorporating empirical functional traits in adaptive landscape analyses requires the continual collection of performance, behavioral, and natural history data across the entire clade.

### Is the carnivoran adaptive landscape relatively flat?

Many carnivoran skeletal components (e.g., mandible, dentition, hindlimb, and post-diaphragmatic region of the vertebral column) exhibit a short phylogenetic half-life relative to the age of Carnivora, suggesting that skeletal traits are strongly pulled towards distinct ecological peaks or clade-based adaptive zones across the adaptive landscape (Slater and Friscia 2019; Law et al. 2022, 2024; Slater 2022). We found that the overall carnivoran landscape based on functional proxies from the skull, appendicular, and axial skeletons can be characterized as a single gradual gradient rather than distinct zones (Fig. 3A). Although some of the performance surfaces and adaptive landscapes show slight ruggedness and multiple peaks and valleys (Fig. 2– 4), we remain cautious in interpreting these as distinct adaptive zones. The use of empirical data in creating performance surfaces and adaptive landscapes can lead to heterogeneous resolution of the interpolation resulting in artificial unevenness in the landscape itself. Our analyses based on theoretical data are more aligned with our views that the topologies of the carnivoran performance surfaces and optimized adaptive landscapes are largely characterized as smooth, gradual gradients with small topographical changes rather than as rugged, multipeak landscapes. Many-to-one mapping (Wainwright et al. 2005) may explain this decoupling between form and function. Multiple combinations of morphological traits may lead to the same functional outcome, resulting in a flat landscape that does not capture the rugged morphological landscape that was previously hypothesized.

The presence of a relatively flat topology may also indicate that carnivorans already occupy a broad adaptive zone relative to the overall mammalian adaptive landscape. Although carnivorans exhibit diverse locomotor modes and correspondingly diverse morphological adaptations, this diversity does not match the extreme locomotor and morphological specialization found in other mammalian clades, especially in the appendicular skeleton such as cranially facing forelimbs in subterranean moles (Lin et al. 2019), digit reduction in cursorial perissodactyls (Economou et al. 2021), and bipedalism in many saltatorial mammals (McGowan and Collins 2018). Many of these specialized mammals may be constrained by their highly derived morphology and thus are adapted to optimize performance for just a single specialized locomotor behavior. That is, most locomotor modes cannot be maximized simultaneously and must trade off with other locomotor modes. In contrast, most carnivorans can perform multiple locomotor behaviors well including running, climbing, digging, and swimming. For example, even the most cursorial carnivoran, the cheetah, can climb trees whereas no cursorial ungulate can. Therefore, the relatively flat landscape in carnivorans signals that functional trade-offs among locomotor performances cannot lead to highly derived specializations, which, in turn, may lead to rugged, multi-peak landscapes in other mammals. Instead, this carnivoran topology highlights that the even slight functional trade-offs across smooth, gradual gradients among appendicular and axial functional proxies can facilitate diverse locomotor modes as well as be flexible enough to enable additional locomotor behaviors. Future work “zooming out” of the carnivoran landscape will further elucidate how the relationships among functional trade-offs, relative degrees of morphological and ecological specializations, and landscape topologies differ among the various clades across Mammalia.

## Acknowledgements

We are grateful to the staff and collections at the American Museum of Natural History, California Academy of Sciences, Field Museum of Natural History, Natural History Museum of Los Angeles County, Museum of Vertebrate Zoology, Natural History Museum London, San Diego Natural History Museum, Texas Vertebrate Paleontology Collection, National Museum of Natural History, and Burke Museum of Natural History and Culture. We thank Blake Dickson and Stephanie Pierce for helpful discussions about adaptive landscapes, Miriam Zelditch and Dave Polly for suggestions on generating theoretical morphological traits from PCAs, and the Santana lab at UW for helpful discussions.

## Authors’ contributions

C.J.L.: conceptualization, data curation, formal analysis, funding acquisition, investigation, methodology, resources, validation, visualization, writing—original draft, writing—review and editing; L.J.H.: funding acquisition, investigation, writing— review and editing; Z.J.T.: funding acquisition, investigation, project administration, writing—review and editing.

