## Supplementary Materials for "The carnivoran adaptive landscape reveals functional trade-offs among the skull, appendicular, and axial skeleton"

Contains:

**Supplementary Methods**

**Fig. S1.** Skeletal trait measurements used in this study.

**Table S1.** PC loadings of skeletal traits.

**Table S3.** Relative weights of functional proxies optimized in the locomotor landscapes.

**Table S4.** Pairwise signiﬁcance tests among dietary adaptive landscapes.

**Table S5.** Relative weights of functional proxies optimized in the dietary landscapes.

**Table S6.** Relative weights of functional proxies optimized in the familial landscapes.

**Table S7.** Pairwise signiﬁcance tests among familial adaptive landscapes.

**Supplementary Results**

**Fig. S2.** Performance surfaces based on theoretical morphology.

**Fig. S3.** Adaptive landscapes optimized for locomotor groups based on theoretical morphology.

**Fig. S4** Adaptive landscapes optimized for dietary groups based on theoretical morphology.

**Fig. S5.** Adaptive landscapes optimized for families based on theoretical morphology.

**Table S8.** Relative weights of functional proxies optimized in the locomotor landscapes.

**Table S9.** Pairwise signiﬁcance tests among locomotor landscapes based on theoretical morphology.

**Table S10.** Pairwise signiﬁcance tests among dietary landscapes based on theoretical morphology.

**Table S11.** Relative weights of functional proxies optimized in the dietary landscapes based on theoretical morphology.

**Table S12.** Relative weights of functional proxies optimized in the familial landscapes based on theoretical morphology.

**Table S13.** Pairwise signiﬁcance tests among familial adaptive landscapes based on theoretical morphology.

**Sensitivity Analyses**

**Fig. S6.** Phylomorphospace of carnivoran skeletal system based on non-size corrected traits.

**Fig. S7.** Performance surfaces from Sensitivity Analyses 4.

**Fig. S8.** Adaptive landscapes from Sensitivity Analyses 4.

**Table S14.** Relative weights of locomotor adaptive landscapes from Sensitivity Analyses 4.

**Table S15.** Relative weights of dietary adaptive landscapes from Sensitivity Analyses 4.

**Table S16.** Relative weights of familial adaptive landscapes from Sensitivity Analyses 4.

**Fig. S9.** Adaptive landscape results from Sensitivity Analyses 2.

**Table S17**. Scaling matrix for the first linear discriminant function.

**Table S18.** Relative weights of landscapes from Sensitivity Analyses 2.

**References**

**Supplementary Methods**

*Creating morphological proxies for function*

From these theoretical morphological traits, we calculated 27 functional indices (Table 1) that capture the functional diversity of the skull (Greaves 2012), limbs (Davis 1964; Samuels et al. 2013), and vertebral column (Boszczyk et al. 2001; Jones et al. 2020, 2021):

Functional proxies for the skull

Temporalis mechanical advantage (temMA) = temporalis moment arm / distance from the jaw condyle to the carnassial. temMA estimates how much force is produced at the bite point from force being input by the temporalis muscle. Lever mechanics suggest that a high temMA indicates a jaw optimized for stronger bites whereas a low temMA indicates a jaw optimized for faster bites. Biomechanical studies have found that increased temMA contributes to increased bite force (Smith and Savage 1956; Throckmorton and Dean 1994; Santana et al. 2012; Law et al. 2016).

Masseter mechanical advantage (masMA) = masseter moment arm / distance from the jaw condyle to the carnassial. masMA estimates how much force is produced at the bite point from force being input by the masseter muscle. Lever mechanics suggest that a high masMA indicates a jaw optimized for stronger bites whereas a low masMA indicates a jaw optimized for faster bites. Biomechanical studies have found that increased masMA contributes to increased bite force (Smith and Savage 1956; Throckmorton and Dean 1994; Santana et al. 2012; Law et al. 2016).

Functional proxies for the forelimb

Scapula index (SI) = scapula length / scapula height. SI describes the expansion of shoulder musculature versus contribution of scapula to limb elongation. The absence or reduction of the clavicle in several mammalian clades allows the scapula to move freely in the parasagittal plane and contribute to stride length (Gasc 2001; Polly 2007; Schmidt and Fischer 2009). Some view the scapula as the first element of the forelimb and thus serves as the functional equivalent to the femur in the hindlimb. Biomechanical analyses found that the proximal elements of the limb (i.e., scapula and femur) produce more than half of the propulsive movement of the whole limb at symmetrical gaits (Fischer et al. 2002). Although the relationship between these performance metrics and scapula shape remains to be tested, ecomorphological studies suggest that increased SI helps with increased stride length and thus tends to be associated with cursorial species, whereas decreased SI provides the teres major muscle with more efficient leverage for powerful adduction of the forelimb and thus tends to be associated with fossorial and natatorial species (Polly 2007).

Brachial index (BI) = radius length / humerus length. BI estimates the relative proportions of the proximal and distal elements of the forelimb and serves as an index of the relative distal out-lever length. The relatively longer distal end of the forelimb facilitates increased stride length and decreased moment of inertia of limbs, which in turn decreases the energetic cost of transport (Kram and Taylor 1990; Strang and Steudel 1990; Garland and Janis 1993; Polly 2007; Pontzer 2007a, 2007b; Kilbourne and Hoffman 2013). Performance studies found that increased BI is correlated with faster running speeds (Christiansen 2002). Ecomorphological studies suggest that increased BI tends to be associated with cursorial species whereas decreased BI is associated with arboreal, natatorial, and fossorial species (Davis 1964; Samuels and Valkenburgh 2008; Samuels et al. 2013).

Humeral robustness index (HRI) = mediolateral diameter of humerus / humerus length. HRI estimates the robustness of the humerus and its ability to resist bending and shearing stresses. Performance studies found that increased HRI is correlated with lower stresses at yield, lower elastic moduli, and higher resistance to failure when a bending force is applied (Kemp et al. 2005). Ecomorphological studies suggest that increased HRI tends to be associated with fossorial and natatorial species whereas decreased HRI is associated with cursorial and scansorial species (Samuels and Valkenburgh 2008; Samuels et al. 2013; Kilbourne 2017; Rickman et al. 2023).

Humeral epicondylar index (HEI) = epicondylar breadth of humerus / humerus length. HEI estimates the relative area of the distal end of the humerus available for the origins of the forearm flexors, pronators, and supinators. Ecomorphological studies suggest that increased HEI tends to be associated with increased fossorial behaviors by helping generate power, force, and stability (Davis 1964; Hildebrand 1985; Lessa and Stein 1992; Lagaria and Youlatos 2006; Samuels and Valkenburgh 2008; Samuels et al. 2013; Rose et al. 2014; Rickman et al. 2023). However, the relationship between HEI and sizes of the forearm flexors, pronators, and supinators remains largely untested.

Olecranon length index (OLI) = olecranon process length / ulna length. OLI estimates the relative mechanical advantage of the triceps brachii and dorsoepitrochlearis muscles used in elbow extension. Biomechanical calculations indicates that an enlarged olecranon process facilitates stronger extension and flexion of the elbow and wrist by increasing mechanical advantage of these muscles, which in turn facilitate the ability to generate large forces (Smith and Savage 1956). Ecomorphological studies suggest that increased OLI is associated with increased fossoriality (Davis 1964; Hildebrand 1985; Lessa and Stein 1992; Lagaria and Youlatos 2006; Samuels and Valkenburgh 2008; Samuels et al. 2013; Rose et al. 2014; Rickman et al. 2023).

Ulnar robustness index (URI) = mediolateral diameter of ulna / ulna length. URI estimates the robustness of the ulna and its ability to resist bending and shearing stresses, and relative area available for the origin and insertion of forearm and manus flexors, pronators, and supinators. Performance studies on URI are limited. URI is presumably similar to radial robustness, where increased radial robustness is correlated with lower stresses at yield, lower elastic moduli, and higher resistance to failure when a bending force is applied (Kemp et al. 2005). Ecomorphological studies suggest that increased URI tends to be associated with fossorial and natatorial species whereas decreased URI tends to be associated with cursorial and scansorial species (Samuels and Valkenburgh 2008; Samuels et al. 2013; Kilbourne 2017; Rickman et al. 2023).

Manus proportions index (MAN) = metacarpal 3 length / humerus length. MAN estimates the relative proportions of proximal and distal elements of the forelimb, and relative size of the hand. Performance studies found that increased MAN is correlated with faster running speeds (Christiansen 2002).

Functional proxies for the hindlimb

Crural index (CI) = tibia length / femur length. CI estimates relative proportions of proximal and distal elements of the hind limb. The relative longer distal end of the hindlimb facilitate increased stride length and decreased moment of inertia of limbs, which in turn decreases the energetic cost of transport (Kram and Taylor 1990; Strang and Steudel 1990; Garland and Janis 1993; Polly 2007; Pontzer 2007a, 2007b; Kilbourne and Hoffman 2013). Performance studies found that increased CI is correlated with faster running speeds (Christiansen 2002). Ecomorphological studies suggest that increased CI tends to be associated with cursorial species whereas decreased CI tends to be associated with arboreal, natatorial, and fossorial species and (Davis 1964; Samuels and Valkenburgh 2008; Samuels et al. 2013).

Femoral robustness index (FRI) = anteroposterior diameter of femur / femur length. Estimates robustness of the femur and its ability to resist bending and shearing stresses. Performance studies found that increased FRI is correlated with lower stresses at yield, lower elastic moduli, and higher resistance to failure when a bending force is applied (Kemp et al. 2005). Ecomorphological studies suggest increased FRI tends to be associated with fossorial and natatorial species whereas decreased FRI tends to be associated with cursorial and scansorial species (Samuels and Valkenburgh 2008; Samuels et al. 2013; Kilbourne 2017; Rickman et al. 2023).

Gluteal index (GI) = length of distal extension of the greater trochanter of the femur / femur length. GI estimates the relative mechanical advantage of the gluteal muscles used in retraction of the femur. Ecomorphological studies suggest increased GI tends to be associated with fossorial (Samuels and Valkenburgh 2008). However, empirical performance data investigating the functional advantages of increased GI remains largely untested.

Femoral epicondylar index (FEI) = epicondylar breadth of femur / femur length. FEI estimates the relative area available for the origins of the gastrocnemius and soleus muscles used in extension of the knee and plantar-flexion of the pes. Ecomorphological studies suggest that increased FEI tends to be associated with increased fossorial behaviors by helping generate power, force, and stability (Davis 1964; Hildebrand 1985; Lessa and Stein 1992; Lagaria and Youlatos 2006; Samuels and Valkenburgh 2008; Samuels et al. 2013; Rose et al. 2014; Rickman et al. 2023). However, empirical performance data investigating the relationship between FEI and sizes of the gastrocnemius and soleus muscles remains largely untested.

Tibial robustness index (TRI) = mediolateral diameter of tibia / tibia length. TRI estimates robustness of the tibia and its ability to resist bending and shearing stresses. Performance studies found that increased TRI is correlated with lower stresses at yield, lower elastic moduli, and higher resistance to failure when a bending force is applied (Kemp et al. 2005). Ecomorphological studies suggest increased TRI tends to be associated with fossorial and natatorial species whereas decreased TRI tends to be associated with cursorial and scansorial species (Samuels and Valkenburgh 2008; Samuels et al. 2013; Kilbourne 2017; Rickman et al. 2023).

Pes length index (PES) = metatarsal 3 length / femur length. Estimates relative proportions of proximal and distal elements of the hind limb, and relative size of the hind foot. Performance studies found that increased PES is correlated with faster running speeds (Garland and Janis 1993; Christiansen 2002).

Functional proxies for the vertebrae

Because the carnivoran vertebral column can be divided into three regionalized modules (i.e., cervical, pre-diaphragmatic thoracic, and post-diaphragmatic thoracolumbar) with the diaphragmatic vertebra serving as a transitional vertebra (Martín-Serra et al. 2021), we only included the functional proxies of the fifth cervical vertebra, middle thoracic vertebra, diaphragmatic vertebra, and middle lumbar vertebra as our proxies for these four regions.

Sagittal second moment of area (sSMA) = π/4 * breadth of posterior centrum * height of the posterior centrum^3. sSMA estimates the stiffness in the of the vertebral joint in the sagittal plane. Biomechanical testing using cadaveric bending experiments found a strong correlation (r = 0.95) between sSMA and compliance (inverse of stiffness) (Jones et al. 2020).

Lateral second moment of area (lSMA) = π/4 * height of posterior centrum * breadth of the posterior centrum^3. lSMA estimates the stiffness in the of the vertebral joint in the lateral plane. Biomechanical testing using cadaveric bending experiments found a strong correlation (r = 0.95) between sSMA and compliance (inverse of stiffness) (Jones et al. 2020).

Joint torsional angle (JTA) = 360º – absolute value of 270º – angle between postzygapophyses. JTA estimates the degree of axial torsion of the vertebrae. Biomechanical testing using cadaveric bending experiments found a strong correlation (r = 0.97) between JTA and mobility torsion of the vertebrae (Jones et al. 2020).

Joint verticality (JV) = absolute value of 180º – angle between postzygapophyses. JV estimates the relative importance of sagittal bending versus lateral bending of vertebral joints. Biomechanical testing using cadaveric bending experiments found a strong correlation (r = 0.77) between JV and the ratio of sagittal to lateral mobility of the vertebrae (Jones et al. 2020).

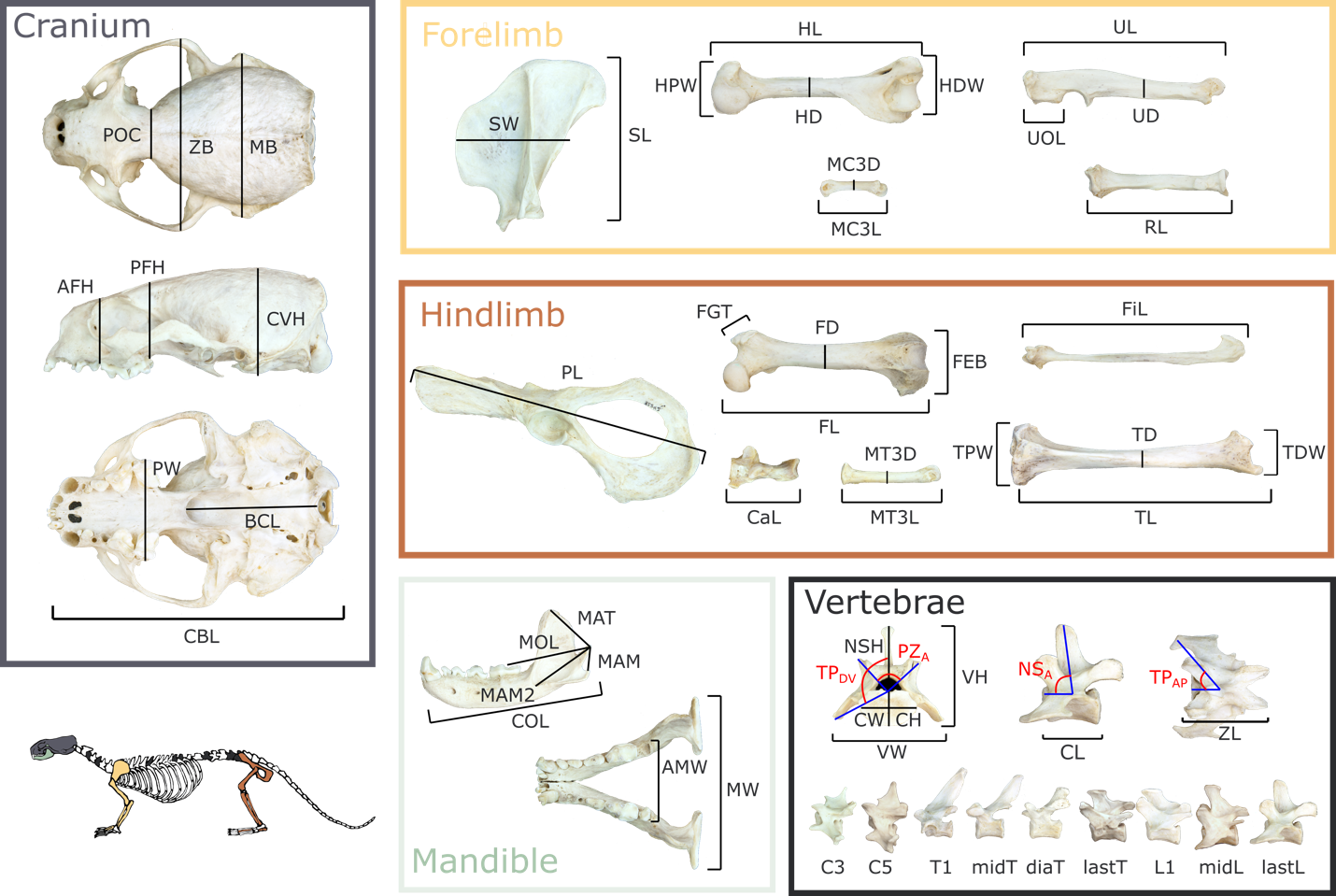

**Fig. S1. Skeletal trait measurements used in this study**. *Cranium*: CBL = condylobasal length, POC = postorbital constriction breadth, ZB = zygomatic breadth, MB = mastoid breadth, AFH = anterior facial height, PFH = posterior facial height, CVH = cranial vault height, PW = palate width, BCL = basicranial length. *Mandible*: MAT = moment arm of temporalis, MAM = moment arm of masseter, MAM2 = moment arm of masseter; MOL = molar out-lever; COL = canine out-lever; MW = mandibular width; AMW = anterior mandibular width. *Forelimb*: SL = scapula length; SW = scapula width; HL = humerus length; HD = humerus mid-shaft width; HPW = humerus proximal width; HDW = humerus distal width; UL = ulna length; UD = ulna mid-shaft width; UOL = ulnar olecranon length; RL = radius length; RD = radius mid-shaft width; MC3L = third metacarpel length; MC3W = third metacarpel width. *Hindlimb*: PL = pelvis length; FL = femur length; FD = femur mid-shaft width; FEB = femur distal width; FGT = height of the greater trochanter of the femur; TL = tibia length; TD = tibia mid-shaft width; TPW = tibia proximal width; TDW = tibia distal width; FiL = fibula length; CaL = calcaneus length; MT3L = third metatarsal length; MT3D = third metatarsal width. *Vertebrae*: VW = vertebrae width; VH = vertebrae height; CL = centrum length; CW = centrum width (posterior); CH = centrum height (posterior); ZL = inter-zygapophyseal length; NSH = neural spine height; TP_DV_ = transverse process dorsoventral projection; TP_AP_ = transverse process anteroposterior projection; PZ_A_ = angle between postzygapophyses; NSA_A_ = neural spine angle.

**Table S1. PC loadings of skeletal traits.** PC 1 and 2 explains 30.3% and 16.5% of the variance, respectively. Trait definitions are found in Figure S1.

| trait | PC1 | PC1 |  | trait | PC1 | PC1 |
| --- | --- | --- | --- | --- | --- | --- |
| lnCBL | 0.050 | -0.058 |  | lnTPW | -0.040 | 0.066 |
| lnPOC | 0.039 | 0.005 |  | lnTDW | -0.053 | 0.050 |
| lnZB | -0.001 | -0.025 |  | lnFiL | 0.130 | 0.118 |
| lnMB | -0.062 | -0.183 |  | lnCaL | 0.033 | 0.100 |
| lnCVH | 0.028 | -0.051 |  | lnMT3L | 0.220 | 0.159 |
| lnPW | 0.074 | -0.048 |  | lnMT3D | 0.038 | 0.123 |
| lnBCL | 0.081 | -0.078 |  | lnC3_VW | -0.045 | -0.087 |
| lnMAT | 0.000 | -0.098 |  | lnC3_VH | 0.042 | -0.026 |
| lnMAM | 0.046 | 0.006 |  | lnC3_CL | 0.137 | 0.054 |
| lnMAM2 | 0.028 | 0.009 |  | lnC3_CW | -0.055 | -0.012 |
| lnMOL | 0.017 | 0.021 |  | lnC3_CH | 0.019 | -0.008 |
| lnCOL | 0.070 | 0.025 |  | lnC3_ZL | 0.118 | 0.021 |
| lnMW | 0.039 | -0.088 |  | lnC3_NSH | 0.057 | -0.039 |
| lnAMW | 0.033 | -0.056 |  | lnC3_TPDV | 0.045 | 0.107 |
| lnSL | 0.007 | 0.143 |  | lnC3_PZA | -0.017 | -0.067 |
| lnSW | -0.056 | 0.111 |  | lnC3_NSA | -0.002 | -0.048 |
| lnHL | 0.049 | 0.110 |  | lnC3_TPAP | 0.040 | 0.020 |
| lnHD | -0.063 | 0.116 |  | lnC5_VW | -0.081 | -0.083 |
| lnHPW | -0.052 | 0.042 |  | lnC5_VH | 0.041 | 0.027 |
| lnHDW | -0.120 | -0.015 |  | lnC5_CL | 0.090 | -0.017 |
| lnUL | 0.053 | 0.206 |  | lnC5_CW | -0.077 | -0.054 |
| lnUD | -0.129 | -0.020 |  | lnC5_CH | 0.034 | 0.054 |
| lnUOL | -0.056 | 0.054 |  | lnC5_ZL | 0.078 | -0.024 |
| lnRL | 0.075 | 0.256 |  | lnC5_NSH | 0.045 | 0.011 |
| lnRD | -0.060 | 0.248 |  | lnC5_TPDV | 0.067 | 0.075 |
| lnMC3L | 0.157 | 0.218 |  | lnC5_PZA | -0.033 | -0.067 |
| lnMC3W | -0.019 | 0.099 |  | lnC5_NSA | -0.044 | -0.095 |
| lnPL | -0.044 | 0.024 |  | lnC5_TPAP | 0.021 | -0.003 |
| lnFL | 0.059 | 0.143 |  | lnT1_VW | -0.067 | -0.062 |
| lnFD | -0.043 | 0.044 |  | lnT1_VH | 0.020 | 0.099 |
| lnFEB | -0.047 | 0.035 |  | lnT1_CL | 0.062 | -0.062 |
| lnFGT | -0.135 | 0.047 |  | lnT1_CW | -0.072 | 0.022 |
| lnTL | 0.124 | 0.114 |  | lnT1_CH | -0.065 | 0.042 |
| lnTD | 0.023 | 0.159 |  | lnT1_ZL | 0.051 | -0.047 |

| trait | PC1 | PC1 |  | trait | PC1 | PC1 |
| --- | --- | --- | --- | --- | --- | --- |
| lnT1_NSH | 0.040 | 0.111 |  | lnlastT_NSA | -0.147 | 0.094 |
| lnT1_TPDV | 0.165 | 0.051 |  | lnL1_VW | -0.085 | 0.077 |
| lnT1_PZA | -0.057 | -0.035 |  | lnL1_VH | -0.106 | 0.088 |
| lnT1_NSA | -0.016 | -0.010 |  | lnL1_CL | 0.093 | -0.079 |
| lnT1_TPAP | -0.040 | 0.027 |  | lnL1_CW | -0.085 | -0.003 |
| lnmidT_VW | -0.052 | -0.050 |  | lnL1_CH | -0.110 | 0.031 |
| lnmidT_VH | -0.040 | 0.129 |  | lnL1_ZL | 0.055 | -0.072 |
| lnmidT_CL | 0.044 | -0.120 |  | lnL1_NSH | -0.104 | 0.117 |
| lnmidT_CW | -0.053 | 0.010 |  | lnL1_TPDV | 0.124 | 0.130 |
| lnmidT_CH | -0.082 | 0.086 |  | lnL1_PZA | -0.059 | -0.126 |
| lnmidT_ZL | 0.013 | -0.114 |  | lnL1_NSA | -0.160 | 0.095 |
| lnmidT_NSH | -0.027 | 0.142 |  | lnL1_TPAP | -0.119 | 0.054 |
| lnmidT_TPDV | 0.176 | 0.042 |  | lnmidL_VW | -0.069 | 0.104 |
| lnmidT_PZA | -0.118 | -0.024 |  | lnmidL_VH | -0.065 | 0.089 |
| lnmidT_NSA | 0.020 | 0.010 |  | lnmidL_CL | 0.128 | -0.065 |
| lnmidT_TPAP | -0.048 | 0.033 |  | lnmidL_CW | -0.081 | 0.003 |
| lntranT_VW | -0.052 | -0.044 |  | lnmidL_CH | -0.118 | 0.061 |
| lntranT_VH | -0.113 | 0.074 |  | lnmidL_ZL | 0.090 | -0.086 |
| lntranT_CL | 0.058 | -0.091 |  | lnmidL_NSH | -0.042 | 0.102 |
| lntranT_CW | -0.060 | 0.000 |  | lnmidL_TPDV | 0.140 | 0.107 |
| lntranT_CH | -0.103 | 0.064 |  | lnmidL_PZA | -0.071 | -0.137 |
| lntranT_ZL | -0.012 | -0.096 |  | lnmidL_NSA | -0.156 | 0.028 |
| lntranT_NSH | -0.117 | 0.076 |  | lnmidL_TPAP | -0.260 | 0.023 |
| lntranT_TPDV | 0.142 | 0.086 |  | lnlastL_VW | -0.049 | 0.068 |
| lntranT_PZA | -0.089 | -0.119 |  | lnlastL_VH | -0.080 | 0.035 |
| lntranT_NSA | 0.014 | 0.032 |  | lnlastL_CL | 0.079 | -0.078 |
| lnlastT_VW | -0.071 | -0.042 |  | lnlastL_CW | -0.058 | 0.019 |
| lnlastT_VH | -0.105 | 0.091 |  | lnlastL_CH | -0.087 | 0.039 |
| lnlastT_CL | 0.079 | -0.091 |  | lnlastL_ZL | 0.059 | -0.100 |
| lnlastT_CW | -0.081 | -0.001 |  | lnlastL_NSH | -0.070 | 0.040 |
| lnlastT_CH | -0.114 | 0.039 |  | lnlastL_TPDV | 0.155 | -0.054 |
| lnlastT_ZL | 0.046 | -0.076 |  | lnlastL_PZA | 0.009 | 0.057 |
| lnlastT_NSH | -0.101 | 0.118 |  | lnlastL_NSA | -0.130 | 0.011 |
| lnlastT_PZA | -0.055 | -0.150 |  | lnlastL_TPAP | -0.207 | -0.013 |

**Table S2. Pairwise tests among ecological regimes.** Bolded P-values indicate significance.

| Locomotor modes | |  |  |  |  |  |  |
| --- | --- | --- | --- | --- | --- | --- | --- |
|  | p-values | arboreal | cursorial | scansorial | semi-aquatic | semi-fossorial | terrestrial hunter |
|  | cursorial | **0.003** |  |  |  |  |  |
|  | scansorial | 0.517 | **0.012** |  |  |  |  |
|  | semi_aquatic | 0.092 | **0.001** | **0.014** |  |  |  |
|  | semi_fossorial | **0.044** | **0.001** | **0.007** | 0.867 |  |  |
|  | terrestrial hunter | 0.079 | **0.012** | **0.025** | **0.001** | **0.001** |  |
|  | terrestrial non-hunter | 0.358 | 0.183 | 0.673 | 0.130 | 0.057 | 0.090 |
| Dietary ecologies | |  |  |  |  |  |  |
|  | p-values | large prey carnivore | med prey carnivore | small prey carnivore | herbivore | insectivore | omnivore |
|  | med prey carnivore | 0.285 |  |  |  |  |  |
|  | small prey carnivore | 0.175 | 0.207 |  |  |  |  |
|  | herbivore | 0.150 | 0.332 | **0.017** |  |  |  |
|  | insectivore | 0.115 | 0.127 | **0.006** | 0.752 |  |  |
|  | omnivore | 0.739 | 0.195 | **0.007** | 0.249 | 0.202 |  |
|  | piscivore | **0.034** | 0.102 | **0.006** | 0.527 | 0.386 | **0.035** |

**Table S3. Relative weights of each theoretical functional proxy optimized in the combined adaptive landscape across all carnivorans and optimized among the seven locomotor adaptive landscapes.** The values represent the mean weights of the traits calculated from the top 5% of landscapes. Functional proxy definitions are found in Table 1.

| functional proxies | combined | arboreal | cursorial | scansorial | semi-aquatic | semi-fossorial | terrestrial hunter | terrestrial non-hunter |
| --- | --- | --- | --- | --- | --- | --- | --- | --- |
| temMA | 0.00 | 0.00 | 0.00 | 0.00 | 0.00 | 0.00 | 0.00 | 0.00 |
| BI | 0.00 | 0.00 | 0.04 | 0.00 | 0.00 | 0.00 | 0.00 | 0.00 |
| HRI | 0.00 | 0.00 | 0.00 | 0.00 | 0.00 | 0.00 | 0.00 | 0.00 |
| HEI | 0.00 | 0.00 | 0.00 | 0.00 | 0.28 | 0.40 | 0.00 | 0.00 |
| OLI | 0.00 | 0.00 | 0.00 | 0.00 | 0.01 | 0.03 | 0.00 | 0.00 |
| MAN | 0.03 | 0.00 | 0.56 | 0.00 | 0.00 | 0.00 | 0.07 | 0.21 |
| CI | 0.03 | 0.00 | 0.00 | 0.00 | 0.03 | 0.00 | 0.00 | 0.01 |
| FRI | 0.00 | 0.00 | 0.00 | 0.00 | 0.00 | 0.00 | 0.00 | 0.00 |
| FEI | 0.00 | 0.00 | 0.00 | 0.00 | 0.07 | 0.00 | 0.00 | 0.00 |
| PES | 0.00 | 0.00 | 0.00 | 0.00 | 0.01 | 0.00 | 0.00 | 0.01 |
| URI | 0.01 | 0.05 | 0.00 | 0.02 | 0.16 | 0.04 | 0.00 | 0.01 |
| SMA_cervical_ | 0.00 | 0.00 | 0.00 | 0.00 | 0.00 | 0.00 | 0.00 | 0.00 |
| JTA_cervical_ | 0.03 | 0.03 | 0.00 | 0.06 | 0.39 | 0.11 | 0.00 | 0.03 |
| JV_cervical_ | 0.03 | 0.03 | 0.02 | 0.00 | 0.00 | 0.00 | 0.07 | 0.03 |
| SMA_thoracic_ | 0.00 | 0.00 | 0.00 | 0.00 | 0.00 | 0.00 | 0.00 | 0.00 |
| JTA_thoracic_ | 0.00 | 0.00 | 0.00 | 0.00 | 0.00 | 0.00 | 0.00 | 0.01 |
| JV_thoracic_ | 0.41 | 0.18 | 0.06 | 0.29 | 0.01 | 0.04 | 0.45 | 0.04 |
| SMA_dia_ | 0.00 | 0.00 | 0.00 | 0.00 | 0.00 | 0.00 | 0.00 | 0.00 |
| JTA_dia_ | 0.03 | 0.03 | 0.00 | 0.04 | 0.04 | 0.37 | 0.00 | 0.60 |
| JV_dia_ | 0.03 | 0.03 | 0.04 | 0.02 | 0.00 | 0.00 | 0.02 | 0.00 |
| SMA_lumbar_ | 0.00 | 0.00 | 0.00 | 0.00 | 0.00 | 0.00 | 0.00 | 0.00 |
| JTA_lumbar_ | 0.00 | 0.00 | 0.00 | 0.00 | 0.00 | 0.00 | 0.00 | 0.03 |
| JV_lumbar_ | 0.38 | 0.68 | 0.29 | 0.56 | 0.00 | 0.00 | 0.39 | 0.03 |
| Z-score | 0.62 | 0.64 | 0.83 | 0.63 | 0.73 | 0.71 | 0.70 | 0.61 |

**Table S4.** **Pairwise signiﬁcance tests among dietary adaptive landscapes.** Top triangle: number of landscape models shared in the top 1% between the paired groups. Bottom triangle: P-values for difference between groups. Bolded P-values indicate significance.

|  | large prey carnivore | med prey carnivore | small prey carnivore | herbivore | insectivore | omnivore | piscivore |
| --- | --- | --- | --- | --- | --- | --- | --- |
| large prey carnivore | - | 4 | 6 | 5 | 5 | 8 | 0 |
| medium prey carnivore | 0.222 | - | 6 | 1 | 8 | 3 | 0 |
| small prey carnivore | 0.333 | 0.462 | - | 3 | 5 | 5 | 0 |
| herbivore | 0.278 | 0.077 | 0.333 | - | 40 | 7 | 1 |
| insectivore | 0.278 | 0.615 | 0.556 | 0.656 | - | 7 | 21 |
| omnivore | 0.444 | 0.231 | 0.556 | 0.115 | **0.045** | - | 0 |
| piscivore | **0.001** | **0.001** | **0.001** | **0.016** | 0.135 | **0.001** | - |

**Table S5.** **Relative weights of each functional proxy optimized among the dietary adaptive landscapes.** The values represent the mean weights of the traits calculated from the top 5% of landscapes. Functional proxy definitions are found in Table 1.

| functional traits | large prey carnivore | med prey carnivore | small prey carnivore | herbivore | insectivore | omnivore | piscivore |
| --- | --- | --- | --- | --- | --- | --- | --- |
| temMA | 0.00 | 0.02 | 0.00 | 0.03 | 0.00 | 0.00 | 0.00 |
| BI | 0.00 | 0.00 | 0.00 | 0.00 | 0.00 | 0.00 | 0.00 |
| HRI | 0.00 | 0.00 | 0.00 | 0.00 | 0.04 | 0.00 | 0.00 |
| HEI | 0.00 | 0.02 | 0.00 | 0.03 | 0.11 | 0.00 | 0.32 |
| OLI | 0.00 | 0.00 | 0.00 | 0.00 | 0.00 | 0.00 | 0.06 |
| MAN | 0.29 | 0.02 | 0.03 | 0.00 | 0.00 | 0.02 | 0.00 |
| CI | 0.00 | 0.00 | 0.00 | 0.00 | 0.00 | 0.02 | 0.03 |
| FRI | 0.00 | 0.00 | 0.00 | 0.00 | 0.00 | 0.00 | 0.00 |
| FEI | 0.00 | 0.00 | 0.00 | 0.00 | 0.00 | 0.00 | 0.11 |
| PES | 0.00 | 0.00 | 0.00 | 0.00 | 0.00 | 0.00 | 0.00 |
| URI | 0.00 | 0.02 | 0.00 | 0.21 | 0.21 | 0.00 | 0.22 |
| SMA_cervical_ | 0.00 | 0.00 | 0.00 | 0.00 | 0.00 | 0.00 | 0.00 |
| JTA_cervical_ | 0.00 | 0.02 | 0.00 | 0.03 | 0.16 | 0.02 | 0.22 |
| JV_cervical_ | 0.01 | 0.04 | 0.03 | 0.03 | 0.00 | 0.02 | 0.00 |
| SMA_thoracic_ | 0.00 | 0.00 | 0.00 | 0.00 | 0.00 | 0.00 | 0.00 |
| JTA_thoracic_ | 0.00 | 0.00 | 0.00 | 0.02 | 0.01 | 0.00 | 0.00 |
| JV_thoracic_ | 0.28 | 0.71 | 0.58 | 0.07 | 0.14 | 0.23 | 0.00 |
| SMA_dia_ | 0.00 | 0.00 | 0.00 | 0.00 | 0.00 | 0.00 | 0.00 |
| JTA_dia_ | 0.00 | 0.04 | 0.00 | 0.20 | 0.19 | 0.02 | 0.03 |
| JV_dia_ | 0.03 | 0.02 | 0.06 | 0.00 | 0.00 | 0.02 | 0.00 |
| SMA_lumbar_ | 0.00 | 0.00 | 0.00 | 0.00 | 0.00 | 0.00 | 0.00 |
| JTA_lumbar_ | 0.00 | 0.00 | 0.00 | 0.00 | 0.01 | 0.00 | 0.00 |
| JV_lumbar_ | 0.39 | 0.10 | 0.31 | 0.38 | 0.13 | 0.64 | 0.00 |
| Z-score | 0.71 | 0.61 | 0.69 | 0.57 | 0.58 | 0.64 | 0.85 |

**Table S6.** **Relative weights of each functional proxy optimized among the carnivoran familial adaptive landscapes.** The values represent the mean weights of the traits calculated from the top 5% of landscapes. Functional proxy definitions are found in Table 1.

| functional traits | combined | Feli | Vive | Hyae | Eupl | Herp | Cani | Ursi | Meph | Proc | Must |
| --- | --- | --- | --- | --- | --- | --- | --- | --- | --- | --- | --- |
| temMA | 0.00 | 0.00 | 0.00 | 0.00 | 0.00 | 0.00 | 0.00 | 0.00 | 0.03 | 0.00 | 0.04 |
| BI | 0.00 | 0.00 | 0.00 | 0.50 | 0.00 | 0.00 | 0.00 | 0.00 | 0.00 | 0.00 | 0.00 |
| HRI | 0.00 | 0.00 | 0.00 | 0.00 | 0.00 | 0.00 | 0.00 | 0.08 | 0.00 | 0.00 | 0.00 |
| HEI | 0.00 | 0.00 | 0.00 | 0.00 | 0.00 | 0.00 | 0.00 | 0.38 | 0.38 | 0.00 | 0.13 |
| OLI | 0.00 | 0.00 | 0.00 | 0.00 | 0.00 | 0.00 | 0.00 | 0.00 | 0.00 | 0.00 | 0.00 |
| MAN | 0.03 | 0.05 | 0.00 | 0.50 | 0.00 | 0.04 | 0.33 | 0.00 | 0.00 | 0.00 | 0.00 |
| CI | 0.03 | 0.00 | 0.00 | 0.00 | 0.04 | 0.04 | 0.00 | 0.00 | 0.03 | 0.00 | 0.04 |
| FRI | 0.00 | 0.00 | 0.00 | 0.00 | 0.00 | 0.00 | 0.00 | 0.00 | 0.00 | 0.00 | 0.00 |
| FEI | 0.00 | 0.00 | 0.00 | 0.00 | 0.00 | 0.00 | 0.00 | 0.00 | 0.00 | 0.00 | 0.02 |
| PES | 0.00 | 0.00 | 0.00 | 0.00 | 0.00 | 0.04 | 0.00 | 0.00 | 0.00 | 0.00 | 0.01 |
| URI | 0.01 | 0.00 | 0.00 | 0.00 | 0.00 | 0.00 | 0.00 | 0.14 | 0.03 | 0.00 | 0.17 |
| SMA_cervical_ | 0.00 | 0.00 | 0.00 | 0.00 | 0.00 | 0.00 | 0.00 | 0.00 | 0.00 | 0.00 | 0.00 |
| JTA_cervical_ | 0.03 | 0.00 | 0.00 | 0.00 | 0.13 | 0.04 | 0.00 | 0.00 | 0.08 | 0.00 | 0.18 |
| JV_cervical_ | 0.03 | 0.02 | 0.07 | 0.00 | 0.00 | 0.00 | 0.05 | 0.00 | 0.00 | 0.00 | 0.00 |
| SMA_thoracic_ | 0.00 | 0.00 | 0.00 | 0.00 | 0.00 | 0.00 | 0.00 | 0.00 | 0.00 | 0.00 | 0.00 |
| JTA_thoracic_ | 0.00 | 0.00 | 0.00 | 0.00 | 0.00 | 0.00 | 0.00 | 0.17 | 0.00 | 0.00 | 0.00 |
| JV_thoracic_ | 0.41 | 0.55 | 0.36 | 0.00 | 0.75 | 0.79 | 0.13 | 0.00 | 0.03 | 0.13 | 0.22 |
| SMA_dia_ | 0.00 | 0.00 | 0.00 | 0.00 | 0.00 | 0.00 | 0.00 | 0.00 | 0.00 | 0.00 | 0.00 |
| JTA_dia_ | 0.03 | 0.00 | 0.00 | 0.00 | 0.04 | 0.04 | 0.00 | 0.24 | 0.38 | 0.00 | 0.13 |
| JV_dia_ | 0.03 | 0.07 | 0.07 | 0.00 | 0.00 | 0.00 | 0.02 | 0.00 | 0.00 | 0.00 | 0.00 |
| SMA_lumbar_ | 0.00 | 0.00 | 0.00 | 0.00 | 0.00 | 0.00 | 0.00 | 0.00 | 0.00 | 0.00 | 0.00 |
| JTA_lumbar_ | 0.00 | 0.00 | 0.00 | 0.00 | 0.00 | 0.04 | 0.00 | 0.00 | 0.00 | 0.00 | 0.00 |
| JV_lumbar_ | 0.38 | 0.32 | 0.50 | 0.00 | 0.04 | 0.00 | 0.47 | 0.00 | 0.03 | 0.88 | 0.05 |
| Z-score | 0.62 | 0.69 | 0.67 | 0.90 | 0.67 | 0.67 | 0.81 | 0.79 | 0.67 | 0.77 | 0.62 |

**Table S7.** **Pairwise signiﬁcance tests among familial adaptive landscapes.** Top triangle: number of landscape models shared in the top 1% between the paired groups. Bottom triangle: P-values for difference between groups. Bolded P-values indicate significance.

|  | Feli | Vive | Hyae | Eupl | Herp | Cani | Ursi | Meph | Proc | Must |
| --- | --- | --- | --- | --- | --- | --- | --- | --- | --- | --- |
| Feli | - | 7 | 0 | 5 | 6 | 35 | 0 | 0 | 8 | 1 |
| Vive | 0.200 | - | 0 | 8 | 10 | 7 | 0 | 0 | 7 | 1 |
| Hyae | **0.001** | **0. 001** | - | 0 | 0 | 0 | 0 | 0 | 0 | 0 |
| Eupl | 0.143 | 0.727 | **0. 001** | - | 8 | 5 | 0 | 0 | 7 | 1 |
| Herp | 0.171 | 0.909 | **0. 001** | 1.000 | - | 6 | 0 | 0 | 7 | 1 |
| Cani | 1.000 | 0.636 | **0. 001** | 0.625 | 0.600 | - | 0 | 0 | 8 | 1 |
| Ursi | **0. 001** | **0. 001** | 0.059 | **0. 001** | **0. 001** | **0. 001** | - | 0 | 0 | 0 |
| Meph | **0. 001** | **0. 001** | **0. 001** | **0. 001** | **0. 001** | **0. 001** | **0. 001** | - | 0 | 58 |
| Proc | 0.229 | 0.636 | **0. 001** | 0.875 | 0.700 | 0.084 | **0. 001** | **0. 001** | - | 4 |
| Must | **0.029** | 0.091 | **0. 001** | 0.125 | 0.100 | **0.011** | **0. 001** | 0.630 | 0.267 | - |

**Supplementary Results**

*Performance surfaces based on theoretical morphology*

The performance surfaces generated based on theoretical morphologies showed similar trends as the performance surfaces generated using the empirical data. In the skull, mechanical advantage of jaw closure tends to be greatest in the bottom of morphospace, with temporalis mechanical advantage (temMA) highest towards the bottom-left (-PC1, -PC2) and declining towards the top-right (+PC1, +PC2) whereas masseter mechanical advantage (masMA) is highest towards the bottom-right (+PC1, -PC2) and declines towards the top-left (-PC1, +PC2) (Fig. S2).

In the forelimb, there is a trade-off between limb elongation and elbow robustness: functional proxies of scapula (SI), radius (BI), and metacarpal (MAN) elongation are highest in the top-right (+PC1, +PC2) of morphospace and transitions towards increased robustness diagonally to the bottom-left (-PC1, -PC2). Proxies associated with increased mechanical advantage of elbow extension (OLI) and attachment sites for forearm flexor, pronator, and supinator muscles on the humeral epicondyles (HEI) and ulna (URI) exhibit the opposite pattern (i.e., highest in the bottom-left and lowest in the top-right). Overall robustness of humerus (HRI) is highest on the left side of the morphospace (-PC1) and transitions towards increased elongation to the right (+PC1), following a similar distribution as the latter indices. The hindlimb also exhibits a trade-off between elongation and robustness: indices of tibial (CI) and metatarsal (PES) elongation tend to be highest in the right side (+PC1, with CI also trending towards -PC2) and transitions to increased robustness towards to the left side (-PC2), whereas indices for femoral (FEI, FRI) and tibial (TRI) robustness tend to be highest on the left side (-PC1) but transitions towards less robustness on PC2 (FEI and FRI is highest towards -PC2 and TRI is highest towards +PC2).

In the vertebral joints, the performance surfaces show a trade-off between joint torsional angle (JTA) as a proxy for range of rotational motion and joint verticality (JV) as a proxy for sagittal mobility. For the cervical, diaphragmatic, and lumbar joints, JTA is highest in the bottom-left of the morphospace (-PC1, -PC2) and declines diagonally to the top-right (+PC1, +PC2), whereas JV exhibits the opposite pattern (i.e., highest in the top-right and lowest in the bottom-left). The thoracic vertebra exhibits similar JTA and JV distributions but in the horizontal plane (i.e., highest JTA in the left side of morphospace and highest JV in the right). In all vertebral joints, second moment of area (SMA) as a proxy for stiffness tends to be greatest towards the top-left of morphospace (-PC1, +PC2) and declines towards the right side of morphospace (+PC1).

*Optimized adaptive landscapes based on theoretical morphology*

After summation of all performance surfaces based on optimized weights, we found that the combined adaptive landscape is heavily weighted for sagittal mobility of the pre-diaphragmatic thoracic region (w_JV_ = 0.71; Fig. S3A; Table S8). When adaptive landscapes are optimized by locomotor ecologies, we found different degrees to which the 27 functional proxies are incorporated among the different adaptive landscapes. The cursorial landscape is characterized by evenly weighted functional proxies associated with increased sagittal mobility of the vertebral column (w_JV_ = 0.13–0.20 for the cervical, pre-diaphragmatic thoracic, diaphragmatic, and lumbar regions) and lengthening of the forelimb, particularly in the scapula (w_SI_ = 0.09), radius (w_BI_ = 0.13), and metacarpal (w_MAN_ = 0.07) (Fig. S3C; Table S8). This landscape is significantly different from all other locomotor landscapes except with the terrestrial non-hunter landscape (Table S9). Semi-aquatic and semi-fossorial landscapes do not significantly differ from one another (P = 0.917; Table S9), and both are weighted for increased joint torsion in the cervical (w_JTA_ = 0.25), diaphragmatic (w_JTA_ = 0.09), and lumbar (w_JTA_ = 0.22) regions; larger femoral epicondyles (w_FEI_ = 0.25); and enlarged olecranon processes (w_OLI_ = 0.09) (Fig. S3E, F; Table S9). The remaining locomotor landscapes do not significantly differ from one another (Table S9) and are heavily weighted for sagittal mobility of the pre-diaphragmatic thoracic region (w_JV_ = 0.51–0.78; Fig. S3; Table S8).

When adaptive landscapes are optimized by dietary ecologies, we found that differences in landscapes appear associated with carnivory (Fig. S4; Table S10). Adaptive landscapes based on diets that largely consists of terrestrial vertebrate prey (i.e., carnivory and omnivory) are heavily weighted for sagittal mobility of the pre-diaphragmatic thoracic region (w_JV_ = 0.66–0.70; Fig. S4; Table S11). In contrast, adaptive landscapes based on diets that do not incorporate terrestrial vertebrate prey (i.e., herbivory, insectivory, and piscivory) are characterized by evenly weighted functional proxies associated with increased joint torsion in the cervical and lumbar regions and larger femoral epicondyles (Fig. S4; Table S11).

Lastly, we found different adaptive landscapes when they are optimized by family. Adaptive landscapes for felids, viverrids, euplerids, herpestids, and procyonids are not significantly different from each other and all remain heavily weighted for sagittal mobility of the pre-diaphragmatic thoracic region (w_JV_ = 0.52–0.79; Fig. S5; Table S12; Table S13). The canid landscape resembles the cursorial landscape with similar weights for increased sagittal mobility of the pre-diaphragmatic thoracic (w_JV_ = 0.36) and diaphragmatic (w_JV_ = 0.23) joints and lengthening of the scapula (w_SI_ = 0.14) (Fig. S5G; Table S12); however, it does not significantly differ from the above family-specific landscapes (Table S13). The mephitid and mustelid landscapes resemble semi-aquatic and semi-fossorial landscapes with similar weights for larger femoral epicondyles (w_FEI_ = 0.21–0.34) and increased joint torsion in the cervical (w_JTA_ = 0.20–0.29) and lumbar (w_JTA_ = 0.16–0.19) joints (Fig. S5I, K), but only the mephitid landscape significantly differs from all other families (Table S13). The hyaenid landscape weighted heavily for increased stiffness of all the cervical joints (w_SMA_ = 0.40) and elongation of the radius (w_BI_ = 0.22) (Fig. S5D), whereas the ursid landscape weighted heavily among increased stiffness of all the vertebral joints (∑w_SMA_ = 0.53) and increased robustness of the humerus (w_HRI_ = 0.18) and tibia (w_TRI_ = 0.20) (Fig. S5H). Both hyaenid and ursid landscapes significantly differ with all other family-specific landscapes (Table S13).

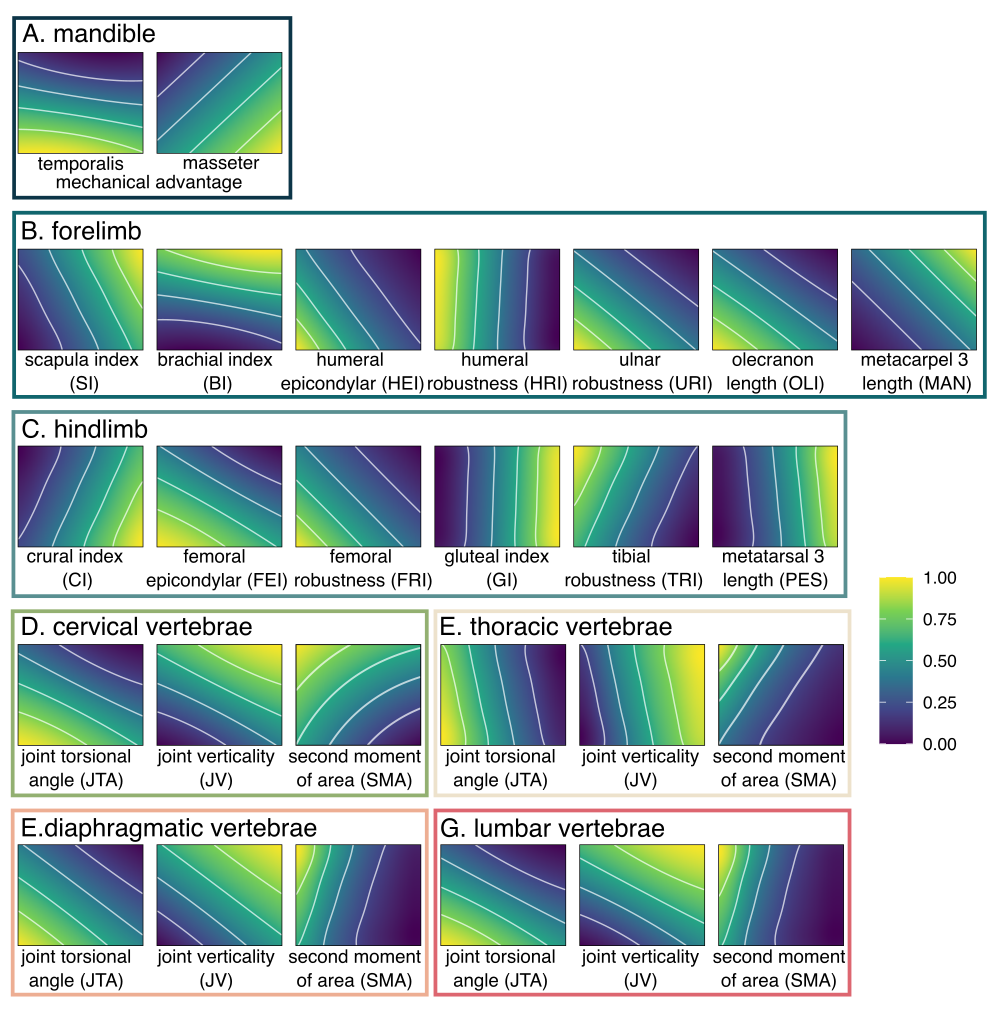

**Fig. S2.** **Performance surfaces for each functional proxy based on theoretical morphology.** Color represents height on the performance surface. See Table 1 for functional proxies definitions.

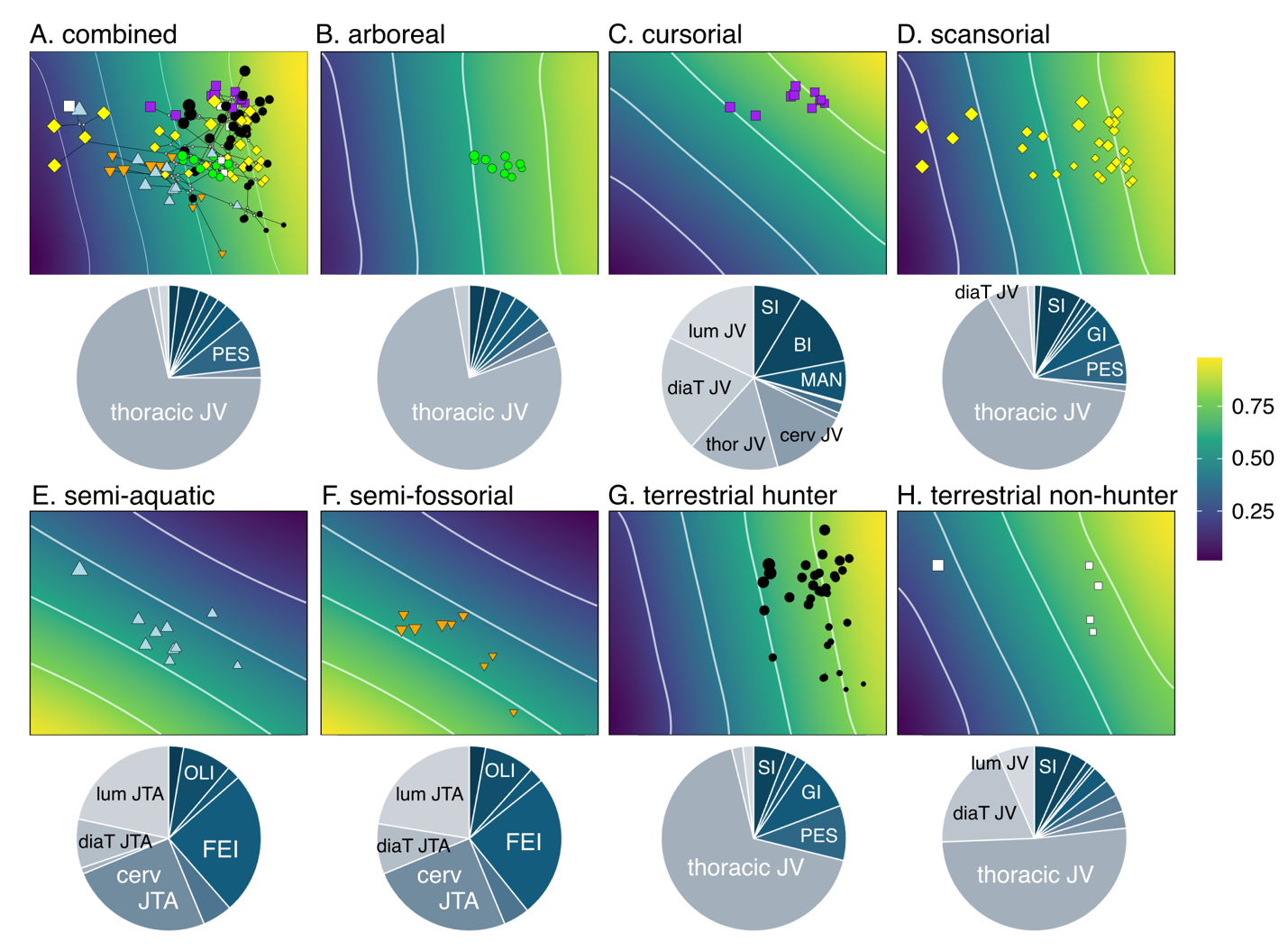

**Fig. S3. Adaptive landscapes optimized for all carnivorans and each locomotor group based on theoretical morphology.** Landscapes were produced by combining the performance surfaces and optimizing their weightings to maximize the height of the landscape at the group mean. Pie charts show the relative weights of each performance surface on each landscape (Table S8 shows breakdown of weights). Functional proxies with weights > 0.07 were labeled. Table S9 shows statistical tests comparing adaptive landscapes among locomotor groups. Size of points are scaled to estimated body size based on the geometric mean of all measurements.

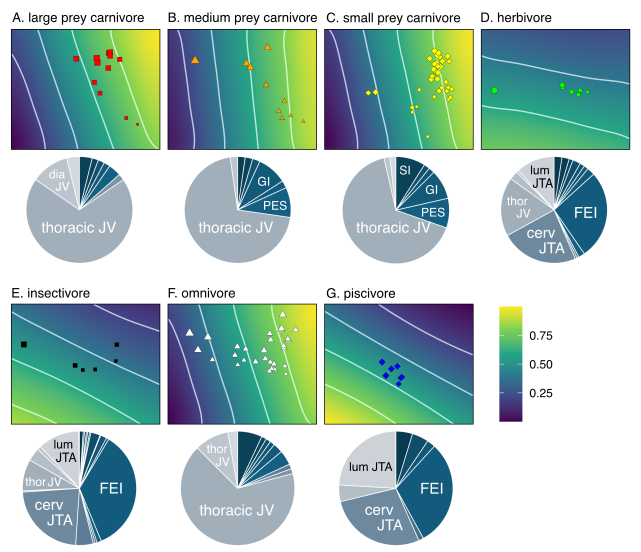

**Fig. S4. Adaptive landscapes optimized for each dietary group based on theoretical morphology.** Adaptive landscapes were produced by combining the performance surfaces and optimizing their weightings to maximize the height of the landscape at the group mean. Pie charts show the relative weighting of each performance surface on each adaptive landscape (see Table S11 for full breakdown of optima weights). Functional indices with weights > 0.09 were labeled. Statistical tests comparing adaptive landscapes among dietary groups can be found in Table S10.

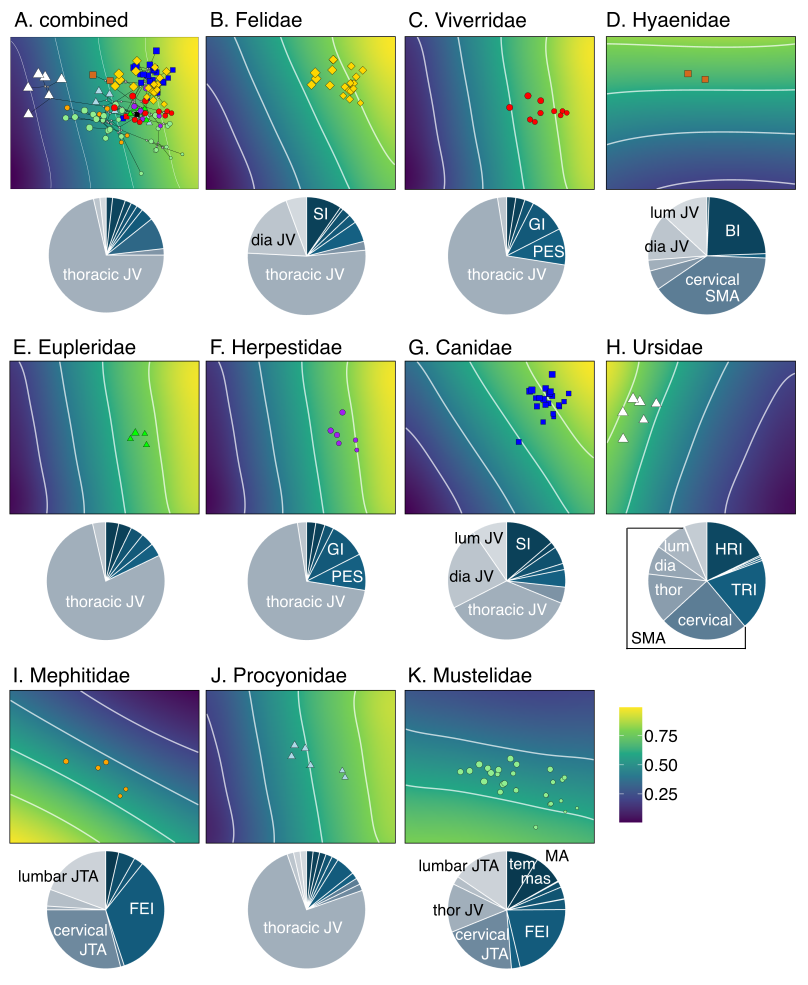

**Fig. S5. Adaptive landscapes optimized for all carnivorans and each family based on theoretical morphology.** Landscapes were produced by combining the performance surfaces and optimizing their weightings to maximize the height of the landscape at the group mean. Pie charts show the relative weights of each performance surface on each adaptive landscape (Table S12 shows breakdown of weights). Functional proxies with weights > 0.09 were labeled. Table S13 shows statistical tests comparing adaptive landscapes among families. Size of points are scaled to estimated body size based on the geometric mean of all measurements.

**Table S8. Relative weights of each theoretical functional proxy optimized in the combined adaptive landscape across all carnivorans and optimized among the seven locomotor adaptive landscapes based on theoretical morphology.** The values represent the mean weights of the traits calculated from the top 5% of landscapes. Functional proxy definitions are found in Table 1.

| functional proxies | combined | arboreal | cursorial | scansorial | semi-aquatic | semi-fossorial | terrestrial hunter | terrestrial non-hunter |
| --- | --- | --- | --- | --- | --- | --- | --- | --- |
| temMA | 0.00 | 0.00 | 0.00 | 0.00 | 0.03 | 0.03 | 0.00 | 0.00 |
| masMA | 0.02 | 0.03 | 0.00 | 0.01 | 0.00 | 0.00 | 0.00 | 0.00 |
| SI | 0.04 | 0.03 | 0.10 | 0.07 | 0.00 | 0.00 | 0.09 | 0.06 |
| BI | 0.02 | 0.00 | 0.15 | 0.01 | 0.00 | 0.00 | 0.00 | 0.07 |
| HRI | 0.00 | 0.00 | 0.00 | 0.00 | 0.00 | 0.00 | 0.00 | 0.00 |
| HEI | 0.00 | 0.00 | 0.00 | 0.00 | 0.00 | 0.00 | 0.00 | 0.00 |
| OLI | 0.00 | 0.00 | 0.00 | 0.00 | 0.09 | 0.09 | 0.00 | 0.00 |
| MAN | 0.02 | 0.00 | 0.10 | 0.01 | 0.00 | 0.00 | 0.02 | 0.02 |
| CI | 0.02 | 0.03 | 0.00 | 0.01 | 0.00 | 0.00 | 0.02 | 0.01 |
| FRI | 0.00 | 0.00 | 0.00 | 0.00 | 0.02 | 0.03 | 0.00 | 0.00 |
| GI | 0.04 | 0.03 | 0.03 | 0.07 | 0.00 | 0.00 | 0.09 | 0.03 |
| FEI | 0.00 | 0.03 | 0.00 | 0.00 | 0.25 | 0.25 | 0.00 | 0.00 |
| TRI | 0.00 | 0.00 | 0.00 | 0.00 | 0.00 | 0.00 | 0.00 | 0.00 |
| PES | 0.09 | 0.03 | 0.05 | 0.08 | 0.00 | 0.00 | 0.09 | 0.05 |
| URI | 0.00 | 0.00 | 0.00 | 0.00 | 0.05 | 0.05 | 0.00 | 0.00 |
| SMA_cervical_ | 0.00 | 0.00 | 0.00 | 0.00 | 0.00 | 0.00 | 0.00 | 0.00 |
| JTA_cervical_ | 0.00 | 0.03 | 0.00 | 0.00 | 0.25 | 0.25 | 0.00 | 0.00 |
| JV_cervical_ | 0.02 | 0.00 | 0.14 | 0.01 | 0.00 | 0.00 | 0.00 | 0.05 |
| SMA_thoracic_ | 0.00 | 0.00 | 0.00 | 0.00 | 0.00 | 0.00 | 0.00 | 0.00 |
| JTA_thoracic_ | 0.00 | 0.00 | 0.00 | 0.00 | 0.00 | 0.00 | 0.00 | 0.00 |
| JV_thoracic_ | 0.71 | 0.78 | 0.14 | 0.67 | 0.01 | 0.00 | 0.66 | 0.56 |
| SMA_dia_ | 0.00 | 0.00 | 0.00 | 0.00 | 0.00 | 0.00 | 0.00 | 0.00 |
| JTA_dia_ | 0.00 | 0.00 | 0.00 | 0.00 | 0.09 | 0.09 | 0.00 | 0.00 |
| JV_dia_ | 0.02 | 0.03 | 0.15 | 0.03 | 0.00 | 0.00 | 0.02 | 0.10 |
| SMA_lumbar_ | 0.00 | 0.00 | 0.00 | 0.00 | 0.00 | 0.00 | 0.00 | 0.00 |
| JTA_lumbar_ | 0.00 | 0.03 | 0.00 | 0.00 | 0.22 | 0.22 | 0.00 | 0.00 |
| JV_lumbar_ | 0.02 | 0.00 | 0.15 | 0.01 | 0.00 | 0.00 | 0.02 | 0.06 |
| Z-score | 0.69 | 0.69 | 0.80 | 0.68 | 0.70 | 0.76 | 0.81 | 0.63 |

**Table S9.** **Pairwise signiﬁcance tests among locomotor adaptive landscapes based on theoretical morphology.** Top triangle: number of landscape models shared in the top 5% between the paired groups. Bottom triangle: P-values for difference between groups. Bolded P-values indicate significance.

|  | arboreal | cursorial | scansorial | semi-aquatic | semi-fossorial | terrestrial hunter | terrestrial non-hunter |
| --- | --- | --- | --- | --- | --- | --- | --- |
| arboreal | - | 4 | 7 | 0 | 0 | 6 | 6 |
| cursorial | 0.400 | - | 10 | 0 | 0 | 7 | 34 |
| scansorial | 0.700 | **0.033** | - | 0 | 0 | 14 | 15 |
| semi-aquatic | **0.001** | **0.001** | **0.001** | - | 88 | 0 | 0 |
| semi-fossorial | **0.001** | **0.001** | **0.001** | 0.917 | - | 0 | 0 |
| terrestrial hunter | 0.600 | **0.023** | 0.778 | **0.001** | **0.001** | - | 12 |
| terrestrial nonhunter | 0.600 | 0.114 | 0.833 | **0.001** | **0.001** | 0.857 | - |

**Table S10.** **Pairwise signiﬁcance tests among dietary adaptive landscapes based on theoretical morphology.** Top triangle: number of landscape models shared in the top 1% between the paired groups. Bottom triangle: P-values for difference between groups. Bolded P-values indicate significance.

|  | large prey carnivore | med prey carnivore | small prey carnivore | herbivore | insectivore | omnivore | piscivore |
| --- | --- | --- | --- | --- | --- | --- | --- |
| large prey carnivore | - | 5 | 7 | 3 | 0 | 11 | 0 |
| med prey carnivore | 0.455 | - | 9 | 7 | 2 | 7 | 0 |
| small prey carnivore | 0.636 | 0.643 | - | 4 | 0 | 11 | 0 |
| herbivore | 0.273 | 0.500 | 0.286 | - | 196 | 4 | 63 |
| insectivore | **0.001** | 0.143 | **0.001** | 0.616 | - | 0 | 62 |
| omnivore | 1.000 | 0.500 | 0.786 | **0.013** | **0.001** | - | 0 |
| piscivore | **0.001** | **0.001** | **0.001** | 0.198 | 0.227 | **0.001** | - |

**Table S11.** **Relative weights of each functional proxy optimized among the dietary adaptive landscapes based on theoretical morphology.** The values represent the mean weights of the traits calculated from the top 5% of landscapes. Functional proxy definitions are found in Table 1.

| functional proxies | large prey carnivore | med prey carnivore | small prey carnivore | herbivore | insectivore | omnivore | piscivore |
| --- | --- | --- | --- | --- | --- | --- | --- |
| temMA | 0.00 | 0.00 | 0.00 | 0.03 | 0.01 | 0.00 | 0.03 |
| masMA | 0.00 | 0.02 | 0.00 | 0.02 | 0.00 | 0.00 | 0.00 |
| SI | 0.05 | 0.02 | 0.09 | 0.01 | 0.00 | 0.08 | 0.00 |
| BI | 0.02 | 0.00 | 0.00 | 0.00 | 0.01 | 0.01 | 0.00 |
| HRI | 0.00 | 0.00 | 0.00 | 0.00 | 0.01 | 0.00 | 0.00 |
| HEI | 0.00 | 0.00 | 0.00 | 0.00 | 0.00 | 0.00 | 0.00 |
| OLI | 0.00 | 0.00 | 0.00 | 0.06 | 0.07 | 0.00 | 0.09 |
| MAN | 0.02 | 0.00 | 0.02 | 0.00 | 0.00 | 0.01 | 0.00 |
| CI | 0.00 | 0.05 | 0.02 | 0.02 | 0.00 | 0.01 | 0.00 |
| FRI | 0.00 | 0.00 | 0.00 | 0.01 | 0.02 | 0.00 | 0.03 |
| GI | 0.02 | 0.09 | 0.09 | 0.02 | 0.01 | 0.03 | 0.00 |
| FEI | 0.00 | 0.02 | 0.00 | 0.20 | 0.23 | 0.00 | 0.25 |
| TRI | 0.00 | 0.00 | 0.00 | 0.00 | 0.01 | 0.00 | 0.00 |
| PES | 0.02 | 0.09 | 0.09 | 0.02 | 0.00 | 0.07 | 0.00 |
| URI | 0.00 | 0.00 | 0.00 | 0.03 | 0.04 | 0.00 | 0.05 |
| SMA_cervical_ | 0.00 | 0.00 | 0.00 | 0.01 | 0.07 | 0.01 | 0.00 |
| JTA_cervical_ | 0.00 | 0.02 | 0.00 | 0.20 | 0.20 | 0.00 | 0.25 |
| JV_cervical_ | 0.02 | 0.00 | 0.00 | 0.00 | 0.00 | 0.01 | 0.00 |
| SMA_thoracic_ | 0.00 | 0.00 | 0.00 | 0.00 | 0.00 | 0.00 | 0.00 |
| JTA_thoracic_ | 0.00 | 0.00 | 0.00 | 0.00 | 0.00 | 0.00 | 0.00 |
| JV_thoracic_ | 0.73 | 0.68 | 0.66 | 0.15 | 0.08 | 0.67 | 0.00 |
| SMA_dia_ | 0.00 | 0.00 | 0.00 | 0.00 | 0.00 | 0.00 | 0.00 |
| JTA_dia_ | 0.00 | 0.00 | 0.00 | 0.06 | 0.07 | 0.00 | 0.09 |
| JV_dia_ | 0.09 | 0.02 | 0.02 | 0.00 | 0.00 | 0.07 | 0.00 |
| SMA_lumbar_ | 0.00 | 0.00 | 0.00 | 0.00 | 0.00 | 0.00 | 0.00 |
| JTA_lumbar_ | 0.00 | 0.00 | 0.00 | 0.16 | 0.15 | 0.00 | 0.22 |
| JV_lumbar_ | 0.02 | 0.00 | 0.02 | 0.00 | 0.00 | 0.01 | 0.00 |
| Z-score | 0.72 | 0.70 | 0.79 | 0.59 | 0.57 | 0.68 | 0.75 |

**Table S12.** **Relative weights of each functional proxy optimized among the carnivoran familial adaptive landscapes based on theoretical morphology.** The values represent the mean weights of the traits calculated from the top 5% of landscapes. Functional proxy definitions are found in Table 1.

| functional proxies | Nand | Feli | Vive | Hyae | Eupl | Herp | Cani | Ursi | Meph | Ailu | Proc | Must |
| --- | --- | --- | --- | --- | --- | --- | --- | --- | --- | --- | --- | --- |
| temMA | 0.00 | 0.00 | 0.00 | 0.00 | 0.00 | 0.00 | 0.00 | 0.00 | 0.03 | 0.02 | 0.00 | 0.08 |
| masMA | 0.04 | 0.00 | 0.02 | 0.00 | 0.03 | 0.03 | 0.00 | 0.00 | 0.00 | 0.02 | 0.02 | 0.06 |
| SI | 0.04 | 0.09 | 0.05 | 0.00 | 0.03 | 0.03 | 0.14 | 0.00 | 0.00 | 0.02 | 0.02 | 0.00 |
| BI | 0.00 | 0.02 | 0.00 | 0.33 | 0.00 | 0.00 | 0.03 | 0.00 | 0.00 | 0.00 | 0.02 | 0.00 |
| HRI | 0.00 | 0.00 | 0.00 | 0.00 | 0.00 | 0.00 | 0.00 | 0.16 | 0.00 | 0.00 | 0.00 | 0.00 |
| HEI | 0.00 | 0.00 | 0.00 | 0.00 | 0.00 | 0.00 | 0.00 | 0.01 | 0.00 | 0.00 | 0.00 | 0.00 |
| OLI | 0.00 | 0.00 | 0.00 | 0.00 | 0.00 | 0.00 | 0.00 | 0.00 | 0.08 | 0.02 | 0.02 | 0.03 |
| MAN | 0.00 | 0.04 | 0.00 | 0.00 | 0.00 | 0.00 | 0.06 | 0.00 | 0.00 | 0.00 | 0.00 | 0.00 |
| CI | 0.04 | 0.00 | 0.02 | 0.00 | 0.03 | 0.03 | 0.00 | 0.00 | 0.00 | 0.02 | 0.02 | 0.03 |
| FRI | 0.00 | 0.00 | 0.00 | 0.00 | 0.00 | 0.00 | 0.00 | 0.01 | 0.02 | 0.00 | 0.00 | 0.00 |
| GI | 0.04 | 0.02 | 0.09 | 0.00 | 0.09 | 0.10 | 0.02 | 0.00 | 0.00 | 0.02 | 0.02 | 0.02 |
| FEI | 0.00 | 0.00 | 0.00 | 0.00 | 0.00 | 0.00 | 0.00 | 0.01 | 0.26 | 0.06 | 0.02 | 0.20 |
| TRI | 0.00 | 0.00 | 0.00 | 0.03 | 0.00 | 0.00 | 0.00 | 0.19 | 0.00 | 0.00 | 0.00 | 0.00 |
| PES | 0.04 | 0.06 | 0.11 | 0.00 | 0.03 | 0.10 | 0.05 | 0.00 | 0.00 | 0.02 | 0.02 | 0.02 |
| URI | 0.00 | 0.00 | 0.00 | 0.00 | 0.00 | 0.00 | 0.00 | 0.00 | 0.04 | 0.00 | 0.00 | 0.01 |
| SMA_cervical_ | 0.00 | 0.00 | 0.00 | 0.39 | 0.00 | 0.00 | 0.00 | 0.24 | 0.00 | 0.00 | 0.02 | 0.00 |
| JTA_cervical_ | 0.00 | 0.00 | 0.00 | 0.00 | 0.00 | 0.00 | 0.00 | 0.00 | 0.26 | 0.06 | 0.02 | 0.22 |
| JV_cervical_ | 0.00 | 0.05 | 0.00 | 0.07 | 0.00 | 0.00 | 0.07 | 0.00 | 0.00 | 0.00 | 0.02 | 0.00 |
| SMA_thoracic_ | 0.00 | 0.00 | 0.00 | 0.00 | 0.00 | 0.00 | 0.00 | 0.13 | 0.00 | 0.00 | 0.00 | 0.00 |
| JTA_thoracic_ | 0.00 | 0.00 | 0.00 | 0.00 | 0.00 | 0.00 | 0.00 | 0.08 | 0.00 | 0.00 | 0.00 | 0.00 |
| JV_thoracic_ | 0.79 | 0.55 | 0.68 | 0.03 | 0.75 | 0.70 | 0.36 | 0.00 | 0.01 | 0.72 | 0.77 | 0.12 |
| SMA_dia_ | 0.00 | 0.00 | 0.00 | 0.00 | 0.00 | 0.00 | 0.00 | 0.10 | 0.00 | 0.00 | 0.00 | 0.00 |
| JTA_dia_ | 0.00 | 0.00 | 0.00 | 0.00 | 0.00 | 0.00 | 0.00 | 0.01 | 0.08 | 0.02 | 0.02 | 0.03 |
| JV_dia_ | 0.04 | 0.11 | 0.02 | 0.05 | 0.03 | 0.03 | 0.17 | 0.00 | 0.00 | 0.02 | 0.02 | 0.00 |
| SMA_lumbar_ | 0.00 | 0.00 | 0.00 | 0.00 | 0.00 | 0.00 | 0.00 | 0.07 | 0.00 | 0.00 | 0.00 | 0.00 |
| JTA_lumbar_ | 0.00 | 0.00 | 0.00 | 0.00 | 0.00 | 0.00 | 0.00 | 0.00 | 0.21 | 0.02 | 0.02 | 0.18 |
| JV_lumbar_ | 0.00 | 0.06 | 0.00 | 0.10 | 0.00 | 0.00 | 0.10 | 0.00 | 0.00 | 0.00 | 0.02 | 0.00 |
| Z-score | 0.73 | 0.81 | 0.78 | 0.69 | 0.74 | 0.78 | 0.83 | 0.89 | 0.70 | 0.66 | 0.62 | 0.68 |

**Table S13.** **Pairwise signiﬁcance tests among familial adaptive landscapes based on theoretical morphology.** Top triangle: number of landscape models shared in the top 1% between the paired groups. Bottom triangle: P-values for difference between groups. Bolded P-values indicate significance.

|  | Nand | Feli | Vive | Hyae | Eupl | Herp | Cani | Ursi | Meph | Ailu | Proc | Must |
| --- | --- | --- | --- | --- | --- | --- | --- | --- | --- | --- | --- | --- |
| Nand | - | 5 | 7 | 0 | 7 | 7 | 5 | 0 | 0 | 7 | 7 | 1 |
| Feli | 0.714 | - | 7 | 0 | 5 | 6 | 35 | 0 | 0 | 5 | 8 | 1 |
| Vive | 1.000 | 0.200 | - | 0 | 8 | 10 | 7 | 0 | 0 | 7 | 7 | 1 |
| Hyae | **0.001** | **0.001** | **0.001** | - | 0 | 0 | 0 | 0 | 0 | 0 | 0 | 0 |
| Eupl | 1.000 | 0.143 | 0.727 | **0.001** | - | 8 | 5 | 0 | 0 | 7 | 7 | 1 |
| Herp | 1.000 | 0.171 | 0.909 | **0.001** | 1.000 | - | 6 | 0 | 0 | 7 | 7 | 1 |
| Cani | 0.714 | 1.000 | 0.636 | **0.001** | 0.625 | 0.600 | - | 0 | 0 | 5 | 8 | 1 |
| Ursi | **0.001** | **0.001** | **0.001** | 0.059 | **0.001** | **0.001** | **0.001** | - | 0 | 0 | 0 | 0 |
| Meph | **0.001** | **0.001** | **0.001** | **0.001** | **0.001** | **0.001** | **0.001** | **0.001** | - | 0 | 0 | 58 |
| Ailu | 1.000 | 0.143 | 0.636 | **0.001** | 0.875 | 0.700 | 0.053 | **0.001** | **0.001** | - | 12 | 8 |
| Proc | 1.000 | 0.229 | 0.636 | **0.001** | 0.875 | 0.700 | 0.084 | **0.001** | **0.001** | 0.750 | - | 4 |
| Must | 0.143 | **0.029** | 0.091 | **0.001** | 0.125 | 0.100 | **0.011** | **0.001** | 0.630 | 0.500 | 0.267 | - |

**Sensitivity Analyses**

Our adaptive landscape results may potentially be affected by (1) removing the effects of size on the morphological traits in creating the morphospace and (2) the coarse increments of partition weights. Therefore, we conducted a series of sensitivity analyses to investigate these potential problems. We used theoretical data to simplify the interpretations of these Sensitivity Analyses.

*Sensitivity Analyses 1: concerns with removing the effects of size in creating the morphospace*

We conducted sensitivity analyses to investigate whether removing the effects of size on the morphological traits in creating the morphospace may potentially influence our performance surfaces and adaptive landscape results. Although it is standard practice to remove the effects of size on morphology analyses, size may be an important predictor of functional performance across Carnivora. Terrestrial carnivorans exhibit a diverse range in body sizes from the 100 g least weasel to 800 kg polar bear. Size effects may therefore facilitate distinct ecological adaptations. We performed a PCA with the same 136 linear and angular measurements but with no size correcting to create the morphospace of the 109 terrestrial carnivoran species. We used the first two PC axes (PC 1 and PC 2 captured 92.4% and 0.02% of the variance, respectively) to create the morphospace (Fig. S6). We then used the same procedure described in the Supplementary Methods to create performance surfaces and optimized adaptive landscapes. We found that functional trade-offs and covariation patterns in the performance surfaces remain similar to the performance surfaces created using size-corrected traits (Fig. S7). The adaptive landscapes optimized by locomotion, diet, and family also exhibit similar topologies and optimized weights as the adaptive landscapes created using size-corrected traits (Fig. S8; Table S14–11).

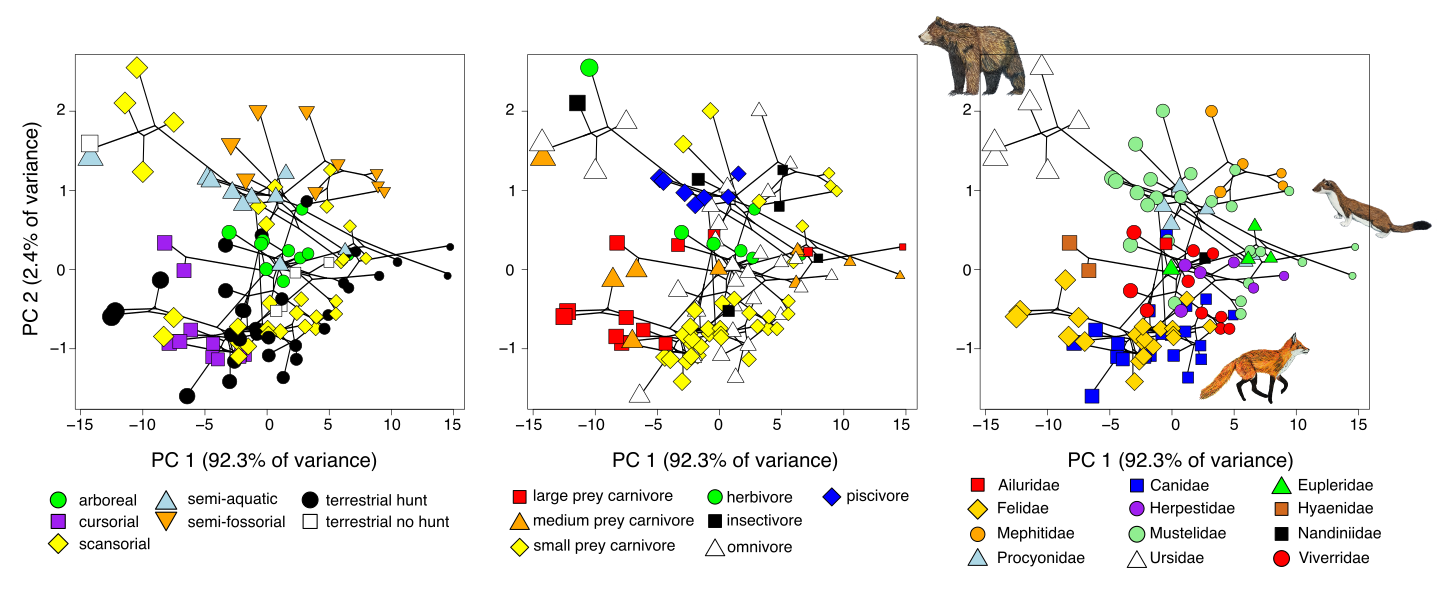

**Fig. S6.** **Phylomorphospace of the carnivoran skeletal system defined by principal components (PCs) 1 and 2.** PCA was conducted using 136 non-size corrected linear and angular measurements that capture morphological variation across the skull, appendicular, and axial skeletons. Size of points are scaled to estimated body size based on the geometric mean of all measurements.

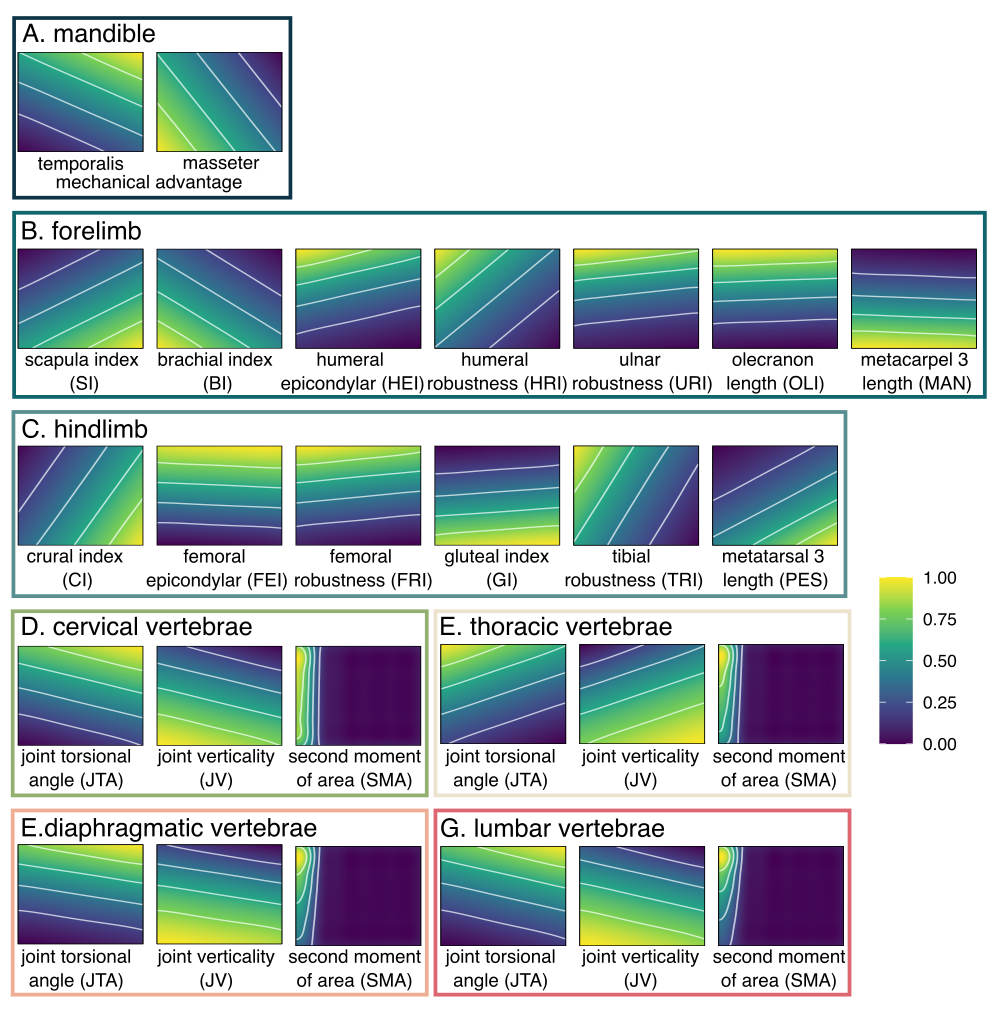

**Fig. S7.** Performance surfaces for each functional proxy based on non-size corrected morphological traits as part of Sensitivity Analyses 4, which investigated whether removing the effects of size on the morphological traits in creating the morphospace may potentially influence our performance surfaces and adaptive landscape results. The largest topological relief correlates with morphospace sample density, suggesting that the unevenness contributes to heterogeneous resolution affect the topology of the performance surfaces. Color represents height on the performance surface.

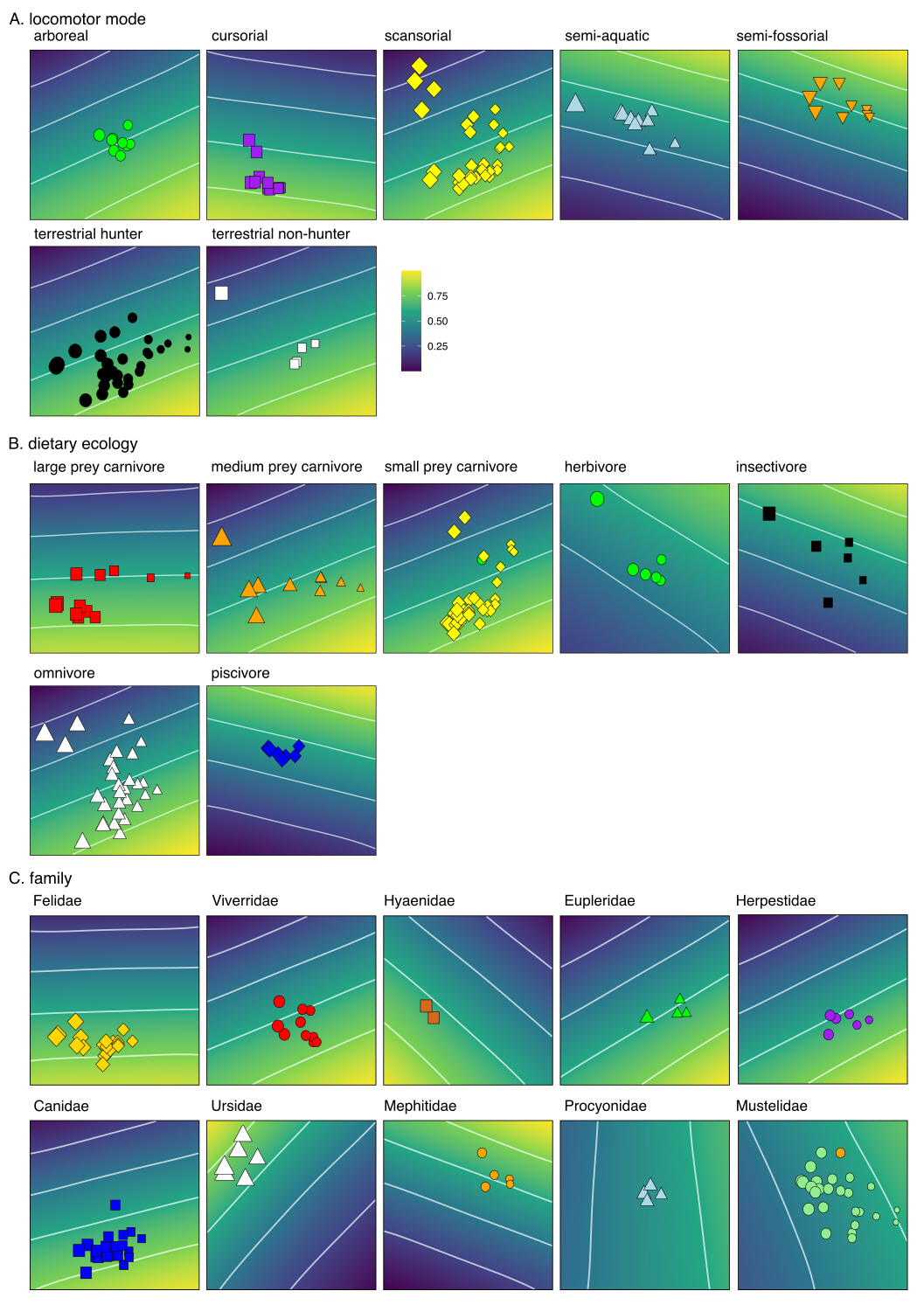

**Fig. S8.** Adaptive landscapes optimized for each locomotor, dietary, and familial groups as part of Sensitivity Analyses 4, which investigated whether removing the effects of size on the morphological traits in creating the morphospace may potentially influence our performance surfaces and adaptive landscape results. Using non-size corrected morphological traits does not change the topologies or weight partitions (Table S14–S11) of the landscapes when optimized for each locomotor group. See Tables S9–S11for breakdown of optima weights. See Figure 3 for original landscapes with all 27 functional proxies.

**Table S14.** Relative weights of each functional proxy optimized among the seven locomotor adaptive landscapes from Sensitivity Analyses 4, which investigated whether removing the effects of size on the morphological traits in creating the morphospace may potentially influence the optimized weights of the adaptive landscape. The values represent the mean weights of the traits calculated from the top 5% of landscapes.

| functional proxies | arboreal | cursorial | scansorial | semi-aquatic | semi-fossorial | terrestrial hunter | terrestrial non-hunter |
| --- | --- | --- | --- | --- | --- | --- | --- |
| temMA | 0.02 | 0.00 | 0.00 | 0.19 | 0.30 | 0.00 | 0.00 |
| masMA | 0.00 | 0.10 | 0.00 | 0.00 | 0.00 | 0.00 | 0.01 |
| SI | 0.09 | 0.05 | 0.18 | 0.00 | 0.00 | 0.21 | 0.13 |
| BI | 0.00 | 0.07 | 0.00 | 0.00 | 0.00 | 0.00 | 0.00 |
| HRI | 0.00 | 0.00 | 0.00 | 0.01 | 0.00 | 0.00 | 0.00 |
| HEI | 0.00 | 0.00 | 0.00 | 0.00 | 0.00 | 0.00 | 0.00 |
| OLI | 0.00 | 0.00 | 0.00 | 0.03 | 0.00 | 0.00 | 0.00 |
| MAN | 0.00 | 0.07 | 0.01 | 0.00 | 0.00 | 0.01 | 0.01 |
| CI | 0.02 | 0.00 | 0.01 | 0.00 | 0.00 | 0.00 | 0.01 |
| FRI | 0.00 | 0.00 | 0.00 | 0.02 | 0.00 | 0.00 | 0.00 |
| GI | 0.02 | 0.11 | 0.03 | 0.00 | 0.00 | 0.04 | 0.03 |
| FEI | 0.00 | 0.00 | 0.00 | 0.15 | 0.07 | 0.00 | 0.00 |
| TRI | 0.00 | 0.00 | 0.00 | 0.03 | 0.00 | 0.00 | 0.00 |
| PES | 0.05 | 0.02 | 0.07 | 0.00 | 0.00 | 0.09 | 0.07 |
| URI | 0.00 | 0.00 | 0.00 | 0.00 | 0.00 | 0.00 | 0.00 |
| SMA_cervical_ | 0.00 | 0.00 | 0.00 | 0.00 | 0.00 | 0.00 | 0.00 |
| JTA_cervical_ | 0.02 | 0.00 | 0.00 | 0.25 | 0.28 | 0.00 | 0.00 |
| JV_cervical_ | 0.00 | 0.13 | 0.01 | 0.00 | 0.00 | 0.01 | 0.01 |
| SMA_thoracic_ | 0.00 | 0.00 | 0.00 | 0.00 | 0.00 | 0.00 | 0.00 |
| JTA_thoracic_ | 0.00 | 0.00 | 0.00 | 0.00 | 0.00 | 0.00 | 0.00 |
| JV_thoracic_ | 0.73 | 0.16 | 0.62 | 0.05 | 0.00 | 0.57 | 0.63 |
| SMA_dia_ | 0.00 | 0.00 | 0.00 | 0.00 | 0.00 | 0.00 | 0.00 |
| JTA_dia_ | 0.00 | 0.00 | 0.00 | 0.10 | 0.10 | 0.00 | 0.00 |
| JV_dia_ | 0.02 | 0.16 | 0.03 | 0.00 | 0.00 | 0.04 | 0.07 |
| SMA_lumbar_ | 0.00 | 0.00 | 0.00 | 0.00 | 0.00 | 0.00 | 0.00 |
| JTA_lumbar_ | 0.00 | 0.00 | 0.00 | 0.16 | 0.25 | 0.00 | 0.00 |
| JV_lumbar_ | 0.02 | 0.15 | 0.03 | 0.00 | 0.00 | 0.03 | 0.03 |
| Z-score | 0.68 | 0.80 | 0.69 | 0.60 | 0.82 | 0.81 | 0.65 |

**Table S15.** Relative weights of each functional proxy optimized among the seven dietary adaptive landscapes from Sensitivity Analyses 4, which investigated whether removing the effects of size on the morphological traits in creating the morphospace may potentially influence the optimized weights of the adaptive landscape. The values represent the mean weights of the traits calculated from the top 5% of landscapes.

| functional proxies | large prey carnivore | med prey carnivore | small prey carnivore | herbivore | insectivore | omnivore | piscivore |
| --- | --- | --- | --- | --- | --- | --- | --- |
| temMA | 0.00 | 0.00 | 0.00 | 0.19 | 0.26 | 0.00 | 0.19 |
| masMA | 0.08 | 0.00 | 0.00 | 0.00 | 0.00 | 0.00 | 0.00 |
| SI | 0.04 | 0.15 | 0.22 | 0.06 | 0.02 | 0.17 | 0.00 |
| BI | 0.02 | 0.00 | 0.00 | 0.00 | 0.00 | 0.00 | 0.00 |
| HRI | 0.00 | 0.00 | 0.00 | 0.00 | 0.00 | 0.00 | 0.00 |
| HEI | 0.00 | 0.00 | 0.00 | 0.00 | 0.00 | 0.00 | 0.00 |
| OLI | 0.00 | 0.00 | 0.00 | 0.02 | 0.02 | 0.00 | 0.03 |
| MAN | 0.02 | 0.00 | 0.01 | 0.00 | 0.00 | 0.02 | 0.00 |
| CI | 0.00 | 0.02 | 0.00 | 0.02 | 0.01 | 0.02 | 0.00 |
| FRI | 0.00 | 0.00 | 0.00 | 0.01 | 0.01 | 0.00 | 0.03 |
| GI | 0.04 | 0.02 | 0.04 | 0.01 | 0.00 | 0.03 | 0.00 |
| FEI | 0.00 | 0.00 | 0.00 | 0.09 | 0.10 | 0.00 | 0.15 |
| TRI | 0.00 | 0.00 | 0.00 | 0.01 | 0.00 | 0.00 | 0.01 |
| PES | 0.01 | 0.08 | 0.09 | 0.03 | 0.01 | 0.08 | 0.00 |
| URI | 0.00 | 0.00 | 0.00 | 0.00 | 0.00 | 0.00 | 0.00 |
| SMA_cervical_ | 0.00 | 0.00 | 0.00 | 0.00 | 0.00 | 0.00 | 0.00 |
| JTA_cervical_ | 0.00 | 0.00 | 0.00 | 0.19 | 0.25 | 0.00 | 0.27 |
| JV_cervical_ | 0.06 | 0.02 | 0.01 | 0.00 | 0.00 | 0.02 | 0.00 |
| SMA_thoracic_ | 0.00 | 0.00 | 0.00 | 0.00 | 0.00 | 0.00 | 0.00 |
| JTA_thoracic_ | 0.00 | 0.00 | 0.00 | 0.00 | 0.00 | 0.00 | 0.00 |
| JV_thoracic_ | 0.43 | 0.65 | 0.57 | 0.16 | 0.06 | 0.63 | 0.01 |
| SMA_dia_ | 0.00 | 0.00 | 0.00 | 0.00 | 0.00 | 0.00 | 0.00 |
| JTA_dia_ | 0.00 | 0.00 | 0.00 | 0.08 | 0.10 | 0.00 | 0.12 |
| JV_dia_ | 0.19 | 0.04 | 0.03 | 0.01 | 0.00 | 0.03 | 0.00 |
| SMA_lumbar_ | 0.00 | 0.00 | 0.00 | 0.00 | 0.00 | 0.00 | 0.00 |
| JTA_lumbar_ | 0.00 | 0.00 | 0.00 | 0.12 | 0.17 | 0.00 | 0.17 |
| JV_lumbar_ | 0.11 | 0.02 | 0.01 | 0.00 | 0.00 | 0.02 | 0.00 |
| Z-score | 0.72 | 0.70 | 0.79 | 0.57 | 0.63 | 0.69 | 0.64 |

**Table S16.** Relative weights of each functional proxy optimized among the familial adaptive landscapes from Sensitivity Analyses 4, which investigated whether removing the effects of size on the morphological traits in creating the morphospace may potentially influence the optimized weights of the adaptive landscape. The values represent the mean weights of the traits calculated from the top 5% of landscapes.

| functional proxies | Feli | Vive | Hyae | Eupl | Herp | Cani | Ursi | Meph | Proc | Must |
| --- | --- | --- | --- | --- | --- | --- | --- | --- | --- | --- |
| temMA | 0.00 | 0.00 | 0.00 | 0.02 | 0.00 | 0.00 | 0.00 | 0.41 | 0.23 | 0.21 |
| masMA | 0.02 | 0.00 | 0.66 | 0.00 | 0.00 | 0.00 | 0.00 | 0.00 | 0.00 | 0.00 |
| SI | 0.07 | 0.18 | 0.00 | 0.27 | 0.30 | 0.13 | 0.00 | 0.00 | 0.09 | 0.11 |
| BI | 0.02 | 0.00 | 0.02 | 0.00 | 0.00 | 0.01 | 0.00 | 0.00 | 0.00 | 0.00 |
| HRI | 0.00 | 0.00 | 0.00 | 0.00 | 0.00 | 0.00 | 0.23 | 0.00 | 0.00 | 0.00 |
| HEI | 0.00 | 0.00 | 0.00 | 0.00 | 0.00 | 0.00 | 0.02 | 0.00 | 0.00 | 0.00 |
| OLI | 0.00 | 0.00 | 0.00 | 0.00 | 0.00 | 0.00 | 0.02 | 0.00 | 0.00 | 0.00 |
| MAN | 0.04 | 0.00 | 0.00 | 0.00 | 0.00 | 0.03 | 0.00 | 0.00 | 0.00 | 0.00 |
| CI | 0.00 | 0.02 | 0.00 | 0.09 | 0.03 | 0.00 | 0.00 | 0.00 | 0.03 | 0.05 |
| FRI | 0.00 | 0.00 | 0.00 | 0.00 | 0.00 | 0.00 | 0.02 | 0.00 | 0.00 | 0.00 |
| GI | 0.09 | 0.03 | 0.02 | 0.00 | 0.02 | 0.08 | 0.00 | 0.00 | 0.00 | 0.01 |
| FEI | 0.00 | 0.00 | 0.00 | 0.00 | 0.00 | 0.00 | 0.03 | 0.03 | 0.03 | 0.04 |
| TRI | 0.00 | 0.00 | 0.02 | 0.00 | 0.00 | 0.00 | 0.57 | 0.00 | 0.00 | 0.00 |
| PES | 0.02 | 0.08 | 0.00 | 0.13 | 0.13 | 0.06 | 0.00 | 0.00 | 0.03 | 0.06 |
| URI | 0.00 | 0.00 | 0.00 | 0.00 | 0.00 | 0.00 | 0.01 | 0.00 | 0.00 | 0.00 |
| SMA_cervical_ | 0.00 | 0.00 | 0.00 | 0.00 | 0.00 | 0.00 | 0.00 | 0.00 | 0.00 | 0.00 |
| JTA_cervical_ | 0.00 | 0.00 | 0.00 | 0.00 | 0.00 | 0.00 | 0.02 | 0.30 | 0.14 | 0.16 |
| JV_cervical_ | 0.09 | 0.02 | 0.02 | 0.00 | 0.00 | 0.03 | 0.00 | 0.00 | 0.00 | 0.00 |
| SMA_thoracic_ | 0.00 | 0.00 | 0.00 | 0.00 | 0.00 | 0.00 | 0.00 | 0.00 | 0.00 | 0.00 |
| JTA_thoracic_ | 0.00 | 0.00 | 0.00 | 0.00 | 0.00 | 0.00 | 0.03 | 0.00 | 0.00 | 0.00 |
| JV_thoracic_ | 0.31 | 0.62 | 0.09 | 0.50 | 0.50 | 0.50 | 0.00 | 0.00 | 0.36 | 0.22 |
| SMA_dia_ | 0.00 | 0.00 | 0.00 | 0.00 | 0.00 | 0.00 | 0.00 | 0.00 | 0.00 | 0.00 |
| JTA_dia_ | 0.00 | 0.00 | 0.00 | 0.00 | 0.00 | 0.00 | 0.02 | 0.05 | 0.02 | 0.05 |
| JV_dia_ | 0.20 | 0.03 | 0.09 | 0.00 | 0.02 | 0.10 | 0.00 | 0.00 | 0.00 | 0.00 |
| SMA_lumbar_ | 0.00 | 0.00 | 0.00 | 0.00 | 0.00 | 0.00 | 0.00 | 0.00 | 0.00 | 0.00 |
| JTA_lumbar_ | 0.00 | 0.00 | 0.00 | 0.00 | 0.00 | 0.00 | 0.02 | 0.21 | 0.06 | 0.10 |
| JV_lumbar_ | 0.15 | 0.02 | 0.09 | 0.00 | 0.00 | 0.06 | 0.00 | 0.00 | 0.00 | 0.00 |
| Z-score | 0.81 | 0.77 | 0.69 | 0.73 | 0.78 | 0.85 | 0.88 | 0.85 | 0.62 | 0.62 |

*Sensitivity Analyses 2: concern with number of partition weight increments*

We conducted sensitivity analyses to investigate whether the coarse increments of partition weights may potentially influence the biological interpretation of our adaptive landscape results. We reduced the number of functional indices by using linear discriminant analyses (LDA) to determine which set of functional proxies best separated out the species among locomotor groups. We conducted one LDA for the limb dataset and a second LDA for the vertebral dataset (see Table S17 for linear discriminant functions). We used LD1 (accounts for 44% of the variance in the limb dataset and 54%% of the variance in the vertebral dataset) to determine the top functional proxies to retain. In the first analysis, we reduced the number of functional proxies to 14 by keeping both mandibular functional proxies (temMA and masMA), the top six vertebral functional proxies (JTA and JV of the cervical, thoracic, and diaphragmatic vertebrae), and the top six limb functional proxies (HRI, HEI, OLI, FRI, FEI, and TRI). This enabled us to reduce the weight partition steps to 0.15, creating a total of 77,520 possible weight combinations. In the second sensitivity analysis, we kept only the top three (JTA and JV of the cervical and JV of the thoracic) vertebral functional proxies and top three limb functional proxies (HRI, HEI, and FRI), enabling us to reduce the weight partition steps to 0.05, creating a total of 53,130 possible weight combinations. We then used the same procedure described in the Supplementary Methods to create performance surfaces and optimized adaptive landscapes. Both sensitivity analyses revealed similar landscape topologies and optimized weights (Fig. S9; Table S18) with the analyses using all 27 functional proxies (Fig. S3). This suggests that the number of functional proxies and subsequent number of steps has minimual impact in determining which traits are heavily weighted on the adaptive landscapes.

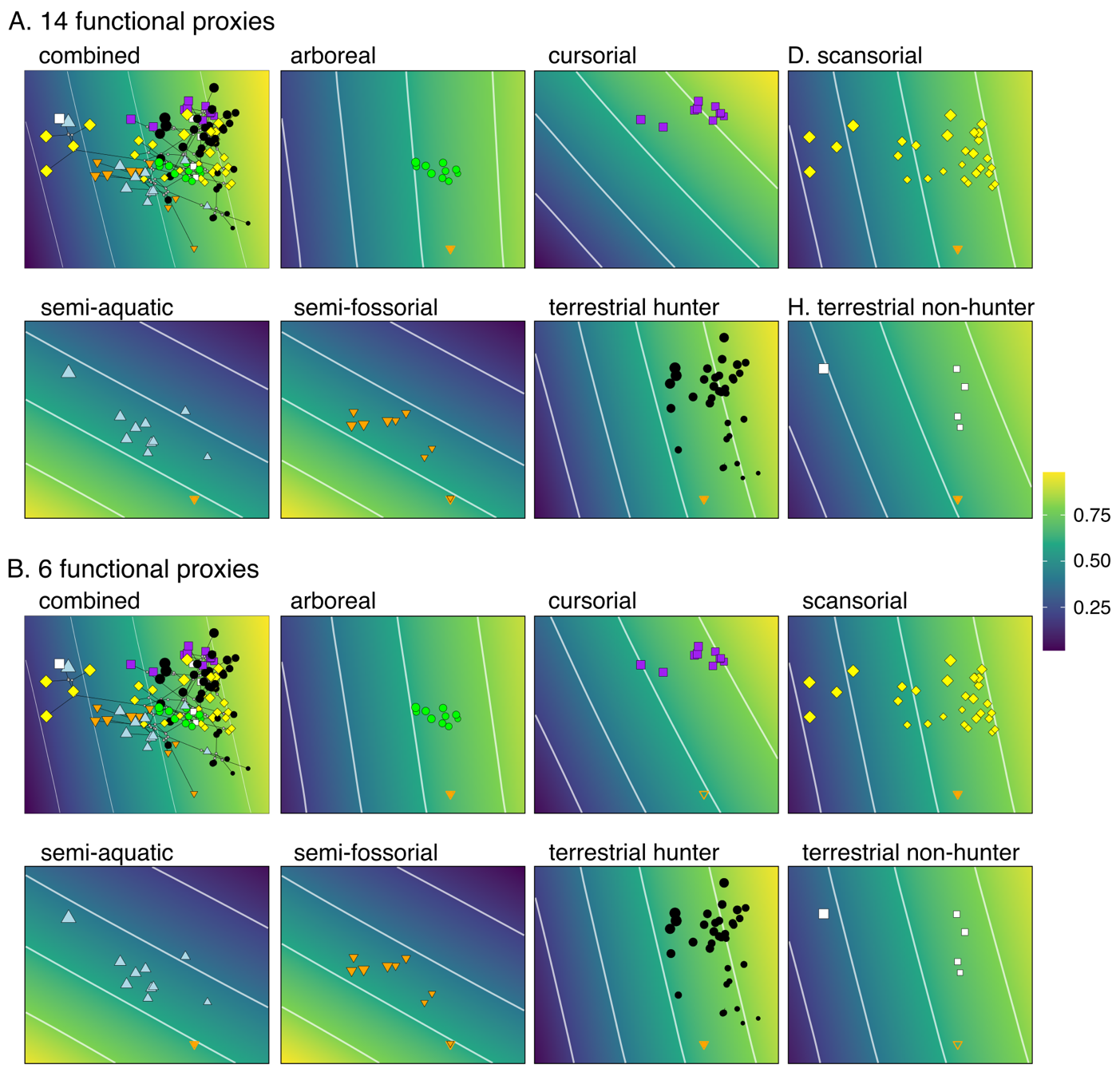

**Fig. S9.** Adaptive landscapes optimized for each locomotor group as part of Sensitivity Analyses 2, which investigated whether the coarse increments of partition weights may potentially influence our adaptive landscape results. Reducing the number of functional proxies from 27 to 14 (weight partition steps reduced to 0.15) or 6 (weight partition steps reduced to 0.07) does not change the topologies or weight partitions (Table S18) of the landscapes when optimized for each locomotor group. See Table S18 for breakdown of optima weights. See Figure 3 for original landscapes with all 27 functional proxies.

**Table S17**. Scaling matrix for the first linear discriminant function (LD1). Separate LDAs were performed for the limb and vertebral datasets.

|  | functional proxies | LD1 |
| --- | --- | --- |
| limbs | |  |
|  | BI | 3.307 |
|  | CI | 3.268 |
|  | FEI | 11.948 |
|  | FRI | 20.308 |
|  | GI | 0.690 |
|  | HEI | 33.182 |
|  | HRI | -83.410 |
|  | IM | 0.474 |
|  | MANUS | -0.506 |
|  | OLI | 10.067 |
|  | PES | 3.282 |
|  | SI | -0.675 |
|  | TRI | 5.948 |
|  | URI | -5.670 |
| vertebrae | |  |
|  | JTA_cervical_ | -0.052 |
|  | JV_cervical_ | 0.052 |
|  | Lateral SMA_cervical_ | 0.000 |
|  | Sagittal SMA_cervical_ | 0.000 |
|  | JTA_lumbar_ | -0.004 |
|  | JV_lumbar_ | 0.004 |
|  | Lateral SMA_lumbar_ | 0.000 |
|  | Sagittal SMA_lumbar_ | 0.000 |
|  | JTA_thoracic_ | -0.013 |
|  | JV_thoracic_ | 0.013 |
|  | Lateral SMA_thoracic_ | 0.000 |
|  | Sagittal SMA_thoracic_ | 0.000 |
|  | JTA_dia_ | -0.013 |
|  | JV_dia_ | 0.013 |
|  | Lateral SMA_dia_ | 0.000 |
|  | Sagittal SMA_dia_ | 0.000 |

**Table S18.** Relative weights of each functional proxy optimized among the seven locomotor adaptive landscapes from Sensitivity Analyses 2, which investigated whether the coarse increments of partition weights may potentially influence our adaptive landscape results. (**A)** Relative weights of proxies when the number of proxies was reduced to 14 and incremental weight partitions was reduced to 0.15. (**B)** Relative weights of proxies when the number of proxies was reduced to 6 and incremental weight partitions was reduced to 0.07. The values represent the mean weights of the traits calculated from the top 5% of landscapes.

|  | functional proxies | arboreal | cursorial | scansorial | semi-aquatic | semi-fossorial | terrestrial hunter | terrestrial non-hunter |
| --- | --- | --- | --- | --- | --- | --- | --- | --- |
| A. number of proxies reduced to 14 | | | | | | | | |
|  | temMA | 0.01 | 0.00 | 0.01 | 0.03 | 0.04 | 0.00 | 0.00 |
|  | masMA | 0.04 | 0.00 | 0.02 | 0.01 | 0.01 | 0.03 | 0.01 |
|  | HRI | 0.00 | 0.00 | 0.00 | 0.01 | 0.00 | 0.00 | 0.01 |
|  | HEI | 0.00 | 0.00 | 0.00 | 0.01 | 0.01 | 0.00 | 0.00 |
|  | OLI | 0.01 | 0.00 | 0.01 | 0.05 | 0.06 | 0.00 | 0.00 |
|  | FRI | 0.01 | 0.00 | 0.00 | 0.03 | 0.03 | 0.00 | 0.00 |
|  | FEI | 0.04 | 0.00 | 0.02 | 0.43 | 0.42 | 0.00 | 0.02 |
|  | TRI | 0.00 | 0.00 | 0.00 | 0.01 | 0.00 | 0.00 | 0.01 |
|  | JTA_cervical_ | 0.04 | 0.00 | 0.02 | 0.35 | 0.36 | 0.00 | 0.01 |
|  | JV_cervical_ | 0.01 | 0.33 | 0.02 | 0.00 | 0.00 | 0.03 | 0.06 |
|  | JTA_thoracic_ | 0.00 | 0.00 | 0.00 | 0.01 | 0.01 | 0.00 | 0.00 |
|  | JV_thoracic_ | 0.79 | 0.33 | 0.80 | 0.02 | 0.01 | 0.86 | 0.64 |
|  | JTA_dia_ | 0.01 | 0.00 | 0.01 | 0.06 | 0.06 | 0.00 | 0.01 |
|  | JV_dia_ | 0.04 | 0.33 | 0.10 | 0.00 | 0.00 | 0.09 | 0.21 |
|  | Z | 0.69 | 0.82 | 0.69 | 0.69 | 0.75 | 0.83 | 0.64 |
| B. number of proxies reduced to 6 | | | | | | | | |
|  | HRI | 0.01 | 0.01 | 0.01 | 0.03 | 0.02 | 0.01 | 0.02 |
|  | HEI | 0.01 | 0.01 | 0.01 | 0.04 | 0.04 | 0.01 | 0.02 |
|  | FRI | 0.02 | 0.01 | 0.02 | 0.07 | 0.07 | 0.01 | 0.02 |
|  | JTA_cervical_ | 0.06 | 0.01 | 0.04 | 0.81 | 0.84 | 0.02 | 0.03 |
|  | JV_cervical_ | 0.03 | 0.47 | 0.06 | 0.01 | 0.01 | 0.05 | 0.09 |
|  | JV_thoracic_ | 0.87 | 0.49 | 0.87 | 0.05 | 0.03 | 0.91 | 0.82 |
|  | Z | 0.69 | 0.79 | 0.69 | 0.68 | 0.74 | 0.82 | 0.64 |
